# Time series scRNAseq analysis in mouse and human informs optimization of rapid astrocyte differentiation protocols

**DOI:** 10.1101/2021.12.07.471509

**Authors:** PW Frazel, D Labib, T Fisher, R Brosh, N Pirianian, A Marchildon, JD Boeke, V Fossati, SA Liddelow

## Abstract

Macroglia (astrocytes and oligodendrocytes) are required for normal development and function of the central nervous system, yet many questions remain about their emergence in the brain and spinal cord. Here we used single-cell RNA sequencing (scRNAseq) to analyze over 298,000 cells and nuclei during macroglia differentiation from mouse embryonic and human induced pluripotent stem cells. We computationally identify candidate genes involved in fate specification of glia in both species, and report heterogeneous expression of astrocyte surface markers across differentiating cells. We then used our scRNAseq data to optimize a previous mouse astrocyte differentiation protocol, decreasing the overall protocol length and complexity. Finally, we used multiomic, dual single nuclei (sn)RNAseq/snATACseq analysis to uncover potential genomic regulatory sites mediating glial differentiation. These datasets enable future optimization of glial differentiation protocols and provide insight into human glial differentiation.

## INTRODUCTION

Macroglia, like astrocytes and oligodendrocytes, are among the last cells to emerge during the self-organization of the mammalian brain. Mapping the molecular mechanisms underpinning their development is integral for understanding the complex physiological heterogeneity of glia^1^, and for understanding how developmental processes may go awry in disease states. Small subgroups of glia respond to insults very differently than other cells^2,3^; thus, understanding developmental processes that may inform this specialization is critical^4^. Single cell RNA sequencing (scRNAseq) has enabled identification of rare cells that play key roles in development and disease, which were previously impossible to observe with methods such as bulk sequencing^5^. The widespread implementation of scRNAseq has produced massive developmental cell atlases of mouse neurodevelopment^6-8^, early human developmental stages^9,10^, and has been used to track heterogeneity during the differentiation of stem cells^11,12^. These analyses have identified genes underpinning key moments in neurodevelopment, such as the emergence of the earliest astrocytes during the “gliogenic switch”^7,13^. In mice, this switch to astrocyte production seems to be controlled by a combination of various transcription factors (TFs) including *Nfia*^14,15^, *Nfib*^16,17^, *Sox9*^18-20^, and signaling pathways like *Notch*^21,22^ (reviewed elsewhere^23-26^). Studies have begun to uncover regional specifications to this process^27-29^ in line with a recent focus on identifying brain-area specific astrocyte subtypes^30-33^.

In humans, much less is understood about the mechanisms directing gliogenesis^25,34^ and neurogenesis more broadly. The development of protocols to generate human astrocyte from pluripotent stem cells (PSCs) has enabled the transcriptional characterization of in vitro (differentiated) astrocytes in comparison with acutely isolated human fetal astrocytes, at bulk and single cell level^35,36^. Conversely, mouse astrocytes have been differentiated from mouse PSCs^37-40^ and characterized through bulk RNA sequencing^37,41^, but the time course of this differentiation in vitro has not yet been analyzed with scRNAseq.

Here, we present scRNAseq analysis of intermediate changes in astrocyte differentiated from both mouse embryonic stem cells (mESCs) and human induced pluripotent stem cells (hiPSCs). We analyzed ∼56,000 cells at 3 timepoints from a previously-verified36 human astrocyte differentiation, and further analyzed another ∼129,000 nuclei across 9 distinct hiPSC lines to ensure consistency across differentiations. In parallel, we prepared and analyzed ∼113,000 cells and nuclei from 15 timepoints across multiple mouse astrocyte differentiation protocols^38^. We computationally identified intermediate and precursor cells in both mouse and human differentiation programs, as well as genes likely to inform fate specification for both species. We used insights from this dataset to shorten and simplify the mouse astrocyte differentiation protocol, and then used the shortened protocol to compare the effects of two different astrocyte differentiation paradigms (BMP4/FGF1 versus CNTF) on gene expression. We also used multiomic sequencing (dual single nuclei (sn) RNAseq/snATACseq) to catalog noncoding genomic regions important for the acquisition of a glial fate, via associations between open chromatin signals and gene expression. Importantly, we report that our rapidly emerging mouse astrocytes demonstrate properties of mature astrocytes: transcriptomic and morphological responses to an inflammatory stimulus, and wound repopulation, measured using a scratch assay. Overall, understanding glia fate specification will facilitate efforts to use these models to understand glial gene expression changes in neurodegeneration.

## RESULTS

### Analysis of intermediate stages during in vitro human

#### astrocyte differentiation

To better understand gliogenesis in hiPSC cultures, we leveraged a serum-free differentiation protocol that generates a mixed culture of astrocytes, neurons, and oligodendrocyte lineage cells, mimicking a more physiological environment in which neural and glial cells grow together. We selected two intermediate timepoints, day 30 and day 50, corresponding to the completion of neurogenesis and the beginning of gliogenesis, respectively^36^, (Fig 1A; Fig S1A) to analyze with scRNAseq. We analyzed a total of 56,446 cells from 6 samples: 33,152 cells were collected from day 30 and day 50, with two independent replicates for each timepoint, and these data were integrated with the data from 23,294 cells at the end of the differentiation (d74) sequenced in a previous study (two technical replicates)^36^. To confirm the robustness of our protocol and address potential inter-line variabilities, we performed independent differentiations using 9 iPSC lines from a collection derived from people with Alzheimer’s disease and unaffected controls, generated using the NYSCF automated reprogramming platform^42^. We then compared gene expression across lines at the single-nucleus level (128,830 nuclei total; see Methods). After integration, there were no major differences in gene expression or cell type representation between lines (Fig S2), corroborating the validity of the in vitro human differentiation to investigate gliogenesis. Cells differentiated from each line were also immunostained for cell-type marker genes, with similar staining patters noted across all 9 lines (Fig S2E). Following quality-control processing via the Muscat^43^ software package, 43,506 cells from the time series dataset remained for analysis (Fig S1B,C). We used Harmony^44^ to integrate for data integration (Fig S1D), and then visualized the output using a force-directed graphical dimensional reduction (FLE^45^; Fig 1B,C). Using this approach, we identified clusters of mature neurons, astrocytes, and oligodendrocytes as expected from our protocol (Fig 1C), based on expression of canonical cell-type marker genes (Fig S3).

**Figure 1.**
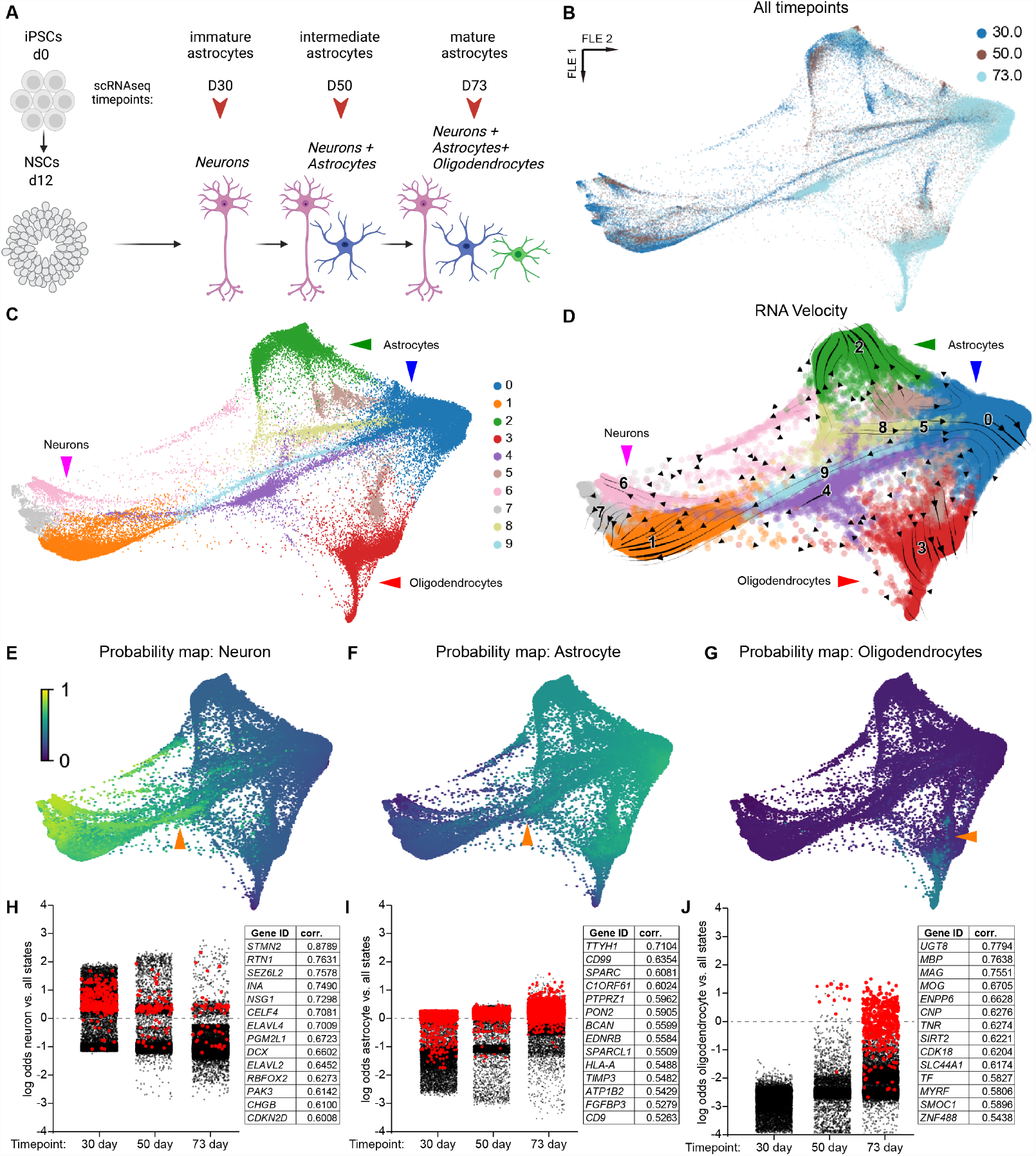
scRNAseq of human glial differentiation uncovers lineage-specific precursors and transient states. A)Human differentiation schematic.B)Dimensional reduction of gene expression for all human data (43,506 cells) shown via force-directed graph (FLE). Each timepoint represents two independent replicates. **C)** FLE graph with cells clustered by gene expression using the Louvain algorithm (resolution: 0.3); mature cell types are identified with arrowheads. **D)** FLE graph with RNA velocity streamlines overlain, to depict flow of cell states over time based on splicing ratios for each gene. **E-G)** Probability maps for each mature cell type based on Waddington Optimal Transport (WOT) algorithm, scale represents the probability for each cell to eventually enter a given state. Orange arrowheads in E-F represent cells likely to become astrocytes or neurons; orange arrowhead in G represents cells likely to become oligodendrocytes. **H-J)** Log odds plots for each mature cell type based on WOT analysis. For every cell from each timepoint, the log odds to become a given cell type versus all other are plotted. In H, red cells express *DCX* > 3 (normalized expression); in I, red cells express *TTYH1* > 3; in J, red cells express *MOG* >3. Each panel also contains a table with the top 15 genes that had expression highly correlated with high probability for a given cell to enter a macrostate (see Methods).

To explore intermediate stages of astrocyte development,we computed RNA velocities^46^ for each cell using the ScVelo^47^ Python software package (Fig 1D). RNA velocity streamlines depict flow of the velocity vectors over time towards more mature clusters (Fig 1D arrowheads; neuron, astrocyte, oligodendrocyte). This temporal ordering is further reinforced by velocity-based pseudotime analysis (Fig S4A), and is consistent with labeling of the samples based on known experimental/differentiation time (Fig 1B). Finally, we observed a clear increase in expression over time for astrocyte marker genes (*CLU, SPARC*) when analyzing all cells per timepoint via pseudobulk (Fig S5B), further underscoring the presence of intermediate glial cells in our dataset.

To better model changes in intermediate cell states we analyzed the data using Waddington Optimal Transport (WOT), an algorithm designed for time-series analysis of scRNAseq data^11^. Implementation of WOT within the CellRank software package^48^ uncovered 4 stable macrostates (Fig S4F), corresponding to two different clusters of mature astrocytes (clusters 0, 2; Fig 1D), and one cluster each of neurons and oligodendrocytes (clusters 6 and 3 respectively; Fig 1D). Marker gene expression for both astrocyte clusters (clusters 0 and 2) are presented in Fig S4H. While cluster 0 expresses standard astrocytic marker genes like *SPARCL1* and *CLU*, cluster 2 marker genes are harder to interpret, suggesting a potential culture artifact (although the top marker gene for Cluster 2 is *SPON1*, a gene enriched in purified human astrocytes). For simplicity, we focused our subsequent analysis on astrocytes from cluster 0, which represent the majority of the astrocytes detected using these methods (15,993 cells in cluster 0; 8,136 cells in cluster 2).

Inspection of the WOT coarse transition matrix confirmed the extreme stability of these 3 macrostates (Fig S4E; see Methods). The CellRank package was then used to calculate the probability for each cell to enter one of the three computationally identified macrostates (astrocytes cluster 0, oligodendrocytes, or neurons). These probability maps suggest that, in vitro, astrocytes and neurons share a common set of early progenitor cells that are likely to enter either state (Fig 1E,F, orange arrowhead; clusters 4 and 9 in Fig 1C,D). These early neural/astrocytic progenitors are also transcriptionally distinct from precursor cells that are likely to become oligodendrocytes (Fig 1G). Cluster 4 potentially represents a multipotent radial-glia-like state, as it persists throughout the differentiation (Fig S3E). Genes enriched in cluster 9 include markers of neurogenesis like *HES6*^49^ and *ASCL1*^50^. This cluster had low cell cycle/proliferation scores (Fig S5B) and decreased in number over time (Fig S3E), suggesting its role as a very early progenitor (i.e. temporally prior to the return to cell cycling that defines early astrogenesis^51^). Consistent with this classification, cells from the proliferative cluster (Fig 1C, cluster 5; see also Fig S6B for proliferation/mitotic scoring) are also very likely to become either astrocytes or oligodendrocytes based on WOT probability maps (Fig 1E-G), but are not likely to become neurons. Finally, these cells also express *NHLH1* (Fig S6C), an understudied TF recently denoted as a neuroblast marker in mouse neurogenesis^7^. In this human dataset, cells that express *NHLH1* are fated to become either neurons or astrocytes, in contrast to the recent finding in mouse^7^.

Based on RNA velocity and fate probability maps, cells from cluster 5 are likely to form both astrocytes and oligodendrocytes, suggesting that this cluster may represent an early group of cells that are committed to gliogenesis. In addition to cell cycle genes like *CENPF* and *MKI67*, enriched genes for this cluster included *HMGB2*, known to be critical for astrogenesis in mice^52^; and published bulk RNA datasets report enrichment for *HMGB2* in acutely purified fetal human astrocytes^53^. We next investigated cells within this cluster that were more likely to become astrocyte versus oligodendrocytes, and similarly attempted to identify putative oligodendrocyte precursor cells (OPCs). Accordingly, the data were re-clustered at a higher resolution (1.2 reclustered versus 0.3 original) to see if cluster 5 would splinter into separate astrocyte- and oligodendrocyte-fated clusters (Fig S6). The resulting cluster (cluster 13, pink arrow) is partially *OLIG1*+ (Fig S6D), but it has low probability of reaching the oligodendrocyte fate (Fig 1G); it is also positive for *OLIG2* (Fig S3C), a key gene in mouse oligodendrogenesis^25^. Thus, these observations are consistent with identifying cluster 13 as a putative pre-OPC state in human cells, potentially similar to a purported pre-OPC state identified recently in a major scRNAseq analysis of mouse neurogenesis^7^. The two clusters directly adjacent to cluster 13 (clusters 4 and 6; green and orange arrows) also have high probability of becoming oligodendrocytes, and express both *EGFR* and *DLL3* (Fig S6C), consistent with identifying these clusters and cluster 13 as pre-OPCs^54^. Pseudobulk analysis of gene expression for these genes reflects this pattern (Fig S7), where expression of OPC genes peaks and then declines during differentiation.

#### Identification of genes correlated with fate decisions

To further investigate intermediate stages in glial development we searched for genes most highly correlated with fate probabilities for a given macrostate. For each cell in the dataset, the odds were computed to reach a given macrostate versus the odds of differentiating into other cell types (Fig 1H-J). Consistent with the known order of emergence of cell types during brain development^7^, the overall odds for cells to become neurons decreased over time (Fig 1H), while the inverse was true for astrocytes and oligodendrocytes (Fig 1I,J). This observation suggests that human CNS cell types are specified in a similar order during both in vitro differentiation and normal development.

To determine how gene expression impacts the likelihood for early cells to differentiate into different types of CNS cells, we plotted cells highly expressing *DCX* (normalized gene expression > 3) in red to trace how decreasing *DCX* expression over the differentiation correlates with decreasing odds to acquire a neuronal cell fate (Fig 1H). Across all three time points, *DCX*-high cells (red) had consistently higher probability of becoming neurons compared to other cells (black). We then used the CellRank package to calculate a Fisher transformation of the correlation coefficient between gene expression and fate probability to identify genes highly correlated with neuronal fate acquisition. *STMN2*, a critical gene for neuron development and maintenance^55^, was most highly correlated with neuronal fate acquisition (r^2^=0.8789). Other genes highly correlated with neuronal fate acquisition included *DCX* (r^2^=0.6602), and two genes implicated in neuron-specific alternative splicing^56^, *ELAVL4* (r^2^=0.7009) and *ELAVL2* (r^2^=0.6452).

When we applied the same analysis to glia, we first found that the gene most correlated with the probability to become an astrocyte was *TTYH1* (r^2^=0.7104), visualized by marking cells with high *TTYH1* expression on the astrocyte-fate log odds plot (normalized expression > 3, Fig 1I). *TTYH1* was not previously implicated in astrocyte differentiation, although recent reports suggested roles for the gene in murine neural stem cell differentiation^57^ and in glioma^58^. Published bulk RNAseq datasets on purified primary astrocytes report *TTYH1* expression is restricted to astrocytes, and increases with age^53^. Interestingly, both *SPARC* and *SPARCL1* expression had similarly strong correlations with astrocyte fate acquisition probability (r^2^=0.6081 and 0.5509 respectively; Fig 1I). This finding supports a recent report^59^ suggesting opposing roles for these two genes during mouse astrocyte development, but with a convergence of mature astrocyte phenotype regardless of the prior *SPARC*/*SPARCL1* status.

Finally, we examined acquisition of oligodendrocyte fate analysis, and identified *UGT8* (r^2^=0.7794) and *MBP* (r^2^=0.7638) as the genes most correlated with oligodendrocyte fate. Accordingly, the log odds graph was labeled for cells highly expressing *MBP* (normalized expression > 3), which marked the cells most likely to become oligodendrocytes at 50 days, and then marked many more cells at 73 days (Fig 1J). This is consistent with the model that oligodendrocytes develop later in differentiation compared to other brain cell types.

Given the important role of TFs in specifying cell fates, we sought to identify TF activities differing between groups of cells. We opted to use BITFAM^60^, a recently developed algorithm for TF activity detection with improved accuracy and speed. Applying BITFAM to the human scRNAseq data revealed multiple TFs with high cluster-specific activity (Fig S8). TFs detected with putative activity in astrocytes included *POU2F2* in the cluster 2 astrocytes, which was recently identified as a potential driver of glioma formation from human astrocytes^61^, in addition to *PAX2*, which plays a key role in astrogenesis in mice^62,63^. Interestingly, in one of the neuron clusters we identified activity of *ARID3B*, a TF understudied in the brain, although one report highlights increased expression in fetal compared to adult mouse brain^64^. *ID1* was highly enriched in two populations of glial progenitors, fitting with its recent identification as an astrocyte-subtype marker in mice^65^. BITFAM also detected *ID3* activity specifically in the oligodendrocyte cluster, even though gene expression of this TF was higher in astrocytes, in line with observations from a recent scRNAseq analysis of human iPSC-derived oligodendrocytes^66^. We also note *NFXL1* activity in the oligodendrocyte cluster, a TF recently associated with a familial speech disorder^67^ and localized to the brain^68^, that has not been previously implicated in oligodendrocyte development.

In sum, we used scRNAseq to analyze differentiating human brain cells in a mixed culture system. Using computational tools we uncovered several genes correlated with cell fate decisions, some of which have not been previously implicated in glial differentiation. These data suggest pathways and genes (e.g., *TTYH1*) that could be targeted during intermediate stages of human glial differentiation to optimize differentiation protocols and/or skew their output towards particular preferred cell types.

#### Optimization of a mouse astrocyte differentiation protocol

Next, we leveraged the insights obtained from the human differentiation analyses to better understand and to optimize the differentiation of astrocytes from mESCs. Much work over the past decade has implicated individual TFs^25^ and signaling molecules in mouse astrocyte differentiation from mESCs, so we sought to exploit the high fidelity afforded by scRNAseq to expand on previous studies. Moreover, a recent report suggested that current murine astrocyte differentiation protocols are unable to produce mature astrocytes^40^, in contrast to the thoroughly verified^36^ human differentiation protocol analyzed above. Other groups have also revealed a similar lack of maturity in astrocytes differentiated without critical developmental cues from neurons^22,37^, underscoring the benefit of the mixed cell-type cultures used in the human dataset, and further emphasizing the need to understand cell fate decisions in order to better optimize murine glial differentiations.

Starting from a serum-free astrocyte differentiation protocol^38^ that produces immature astrocytes from mESCs, we developed a modified protocol aimed at producing CNS astrocytes (Fig 2A; Fig S9A). Modifications initially included: 1) removing CNTF as a growth factor due to recent reports that it induces a reactive state in astrocytes^37^; 2) coating plates with poly-d-lysine and laminin during later protocol stages;and 3) removing retinoic acid and smoothened agonist to avoid induction of spinal cord identity^69,70^. We tested that this modified protocol eventually produces astrocytes by stimulating cells with CNTF and measuring *Gfap* expression, as astrocytes upregulate *Gfap* in response to many reactive signals^2,71^. Robust GFAP staining levels were observed in cells differentiated from two distinct mESC lines after 12 days total using the modified differentiation: 4 days of embryoid body (EB) expansion, 4 days of adherent differentiation, then 4 days of differentiation with CNTF (Fig 2B). This was significantly faster than the onset of *Gfap* expression at 21 days, as noted for the original differentiation protocol^38^, although the modified differentiation also replicated the finding of lower GFAP levels in FGF/BMP4-treated cells (Fig S9E,F). This difference in Gfap expression timing are likely due to protocol modifications discussed above, although additionally, in the original study timepoints prior to 21 days were not analyzed. To ensure that the rapid Gfap induction was not caused by the use of a different mESC line compared to the original protocol38, a second mESC line was differentiated and also displayed robust GFAP immunostaining (Fig 2B). This high level of Gfap after stimulation with CNTF further supports the decision to exclude CNTF (at first) due to concerns of producing reactive cells^2^,^71^.

**Figure 2.**
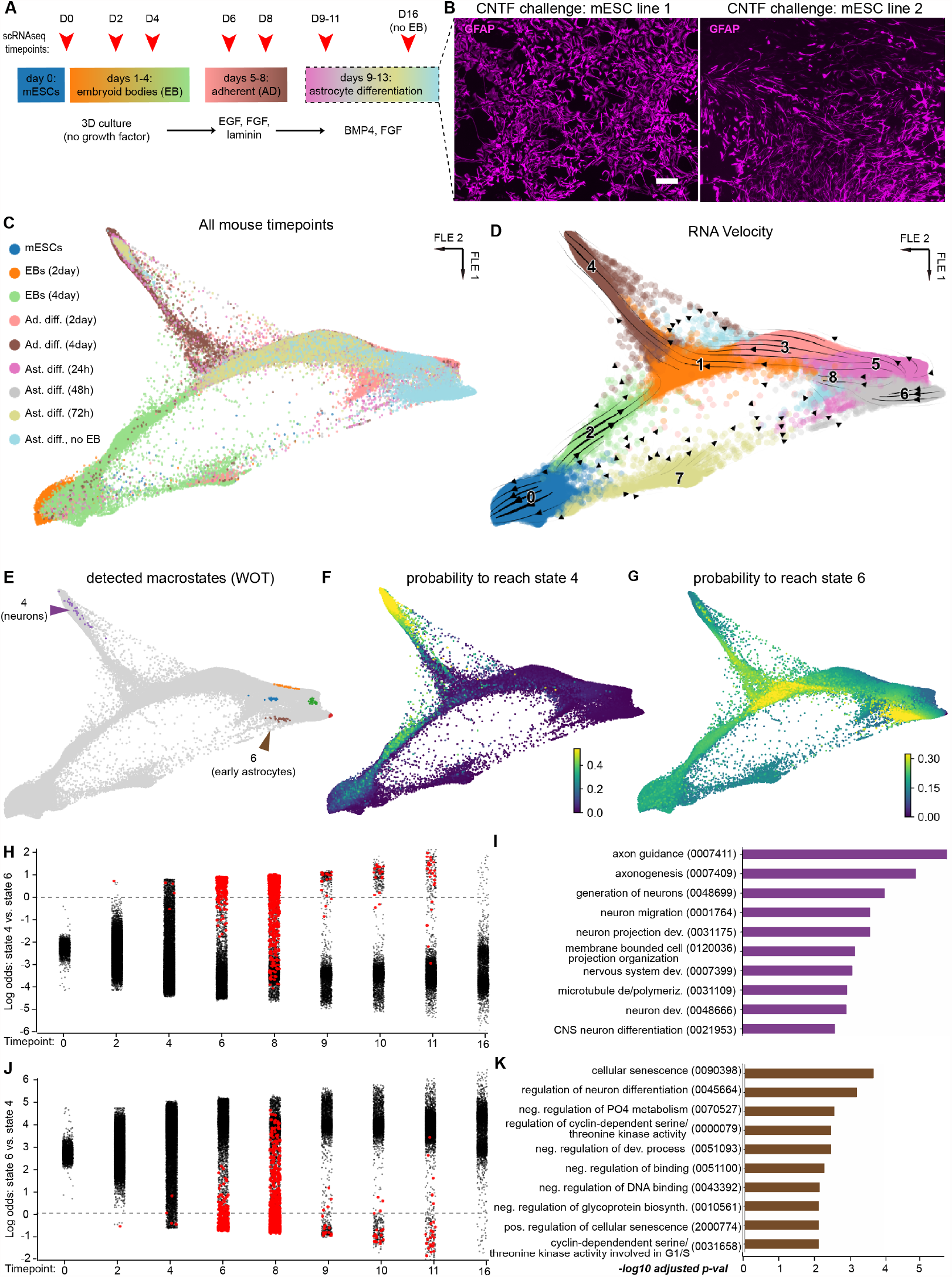
scRNAseq of mouse glial differentiation uncovers lineage-specific precursors and transient states. **A)**Mouse differentiation schematic.**B)**Micrograph of differentiated astrocytes displaying robust GFAP staining after 12 days of differentiation with CNTF (CNTF was not used for any subsequent differentiations in this figure). mESC line 1: A17iCre, mESC line2: Bl6/CAST dPIGA. Scale bar = 100 µm. **C)** Force-directed graph (FLE) graph of all mouse timepoints (58,268 cells total). **D)** FLE graph with RNA velocity streamlines overlain, to depict flow of cell states over time based on splicing ratios for each gene. Cells were clustered with the Louvain algorithm (resolution: 0.7); clusters are labeled by number on the plot. **E)** Multiple macrostates were detected with the Waddington Optimal Transport (WOT) algorithm; arrowheads depict neurons (state 4) and early astrocytes (state 6). **F**,**G)** Probability maps for each mature cell type based on the WOT algorithm, scale represents the probability for each cell to eventually enter a given state. **H**,**J)** Log odds plots for each mature cell type based on WOT analysis. For every cell from each timepoint, the log odds to become a given cell type versus the other cell type are plotted. In both panels, red cells express *DCX* > 1. **I**,**K)** Gene ontology (GO) term analysis of genes identified via WOT as driving fate commitment for neuron (purple) and early astrocyte (brown) macrostates. GO IDs are in parentheses next to their descriptor.

#### Early stages of mouse astrocyte differentiation

To better understand early astrocyte development, we analyzed gene expression during the first 12 differentiation days using scRNAseq, examining a total of 58,268 mouse cells after preprocessing with Muscat for quality control (Fig S9B,G-I). The first mouse dataset spans 9 timepoints (Fig 2A,C): from mESCs to embryoid bodies (EBs), through the intermediate adherent differentiation (AD) stage, and finally through exposure to FGF1/BMP4 (Fig 2A). As with the human data, batch effects were corrected using Harmony (Fig S9G). RNA velocity demonstrated a clear velocity flow away from the mESC and EB stages towards the intermediate stages (AD and FGF1/BMP4 exposure; Fig 2D). Similar to the velocity results in the human dataset (Fig 1D), the velocity vectors for many of the intermediate stages were incoherent, which is a known aspect of velocity analysis^72^. Further, even though computed RNA velocity was high (Fig S10B), and the velocity streamlines aligned with known external timepoints (Fig 2D), the calculated velocity pseudotime was inaccurate based on the known order of the time series (Fig S10A). Calculating latent time based on the dynamic model of RNA velocity also failed to produce an accurate pseudotime (Fig S10C).

To employ non-RNA velocity modeling, the mouse dataset was analyzed with the WOT time series algorithm. The algorithm detected multiple macrostates consistent with the samples’ known temporal order, including one neuronal macrostate (state 4) and several non-neuronal states (Fig 2E). State 4 was assigned as neuronal given its high expression of *Stmn2, Dcx*, and *Tubb3* (Fig S11A), while state 5 is likely an early astrocytic progenitor based on enrichment for genes like *Vim, Sparcl1*, and *Clu* (Fig S11B). Identification of only two macrostates by the WOT algorithm was unsurprising, as this mouse astrocyte differentiation is not expected to produce a third cell type (oligodendrocytes), in contrast to the human protocol (Fig 1 and Discussion).

We next calculated the probability for each cell to enter either of the two macrostates (Fig 2F,G) in addition to genes correlated with probability to enter each state (Fig 2H, I). Consistent with the overall arc of the differentiation towards astrocyte production, most cells in the dataset had a low chance of entering the neuronal state (Fig 2F), and instead had high odds to reach the non-neuronal state (Fig 2G). To investigate how the probability of reaching each state changes over the differentiation, the log odds to reach one state versus the other across all 9 timepoints were plotted and analyzed (Fig 2H, J). There was an overall decrease in the odds of reaching the neuronal state (state 4) over time, consistent with pseudobulk data demonstrating dramatic decreases in overall expression of neurogenic genes like *Dcx, Tubb3*, and *Stmn2* across the span of the differentiation (Fig S12A).

We focused on the relationship between *Dcx* expression and neuronal fate acquisition by marking cells red if they expressed *Dcx* above a normalized threshold of 1 (Fig 2H, J). Mirroring findings from the human dataset, *Dcx* expression was clearly associated with higher odds to reach the neuronal state versus nonneuronal (Fig 2H), and conversely was anticorrelated with a cells’ chances to reach the nonneuronal fate (state 6; Fig 2J). Consistent with these observations, *Dcx* was one of the top 10 genes whose expression was most correlated with odds to reach state 4 (neuronal) versus state 6 (Fig S10I). *Nhlh1* and *Nhlh2* were enriched in cells with high neuronal fate probability (Fig S11C), two TFs recently suggested to mark a neuroblast population that exclusively generates neurons^7^. Finally, gene ontology (GO) analysis of the genes most correlated with neuronal fate (macrostate 4) was enriched for terms associated with neurogenesis (Fig 2I).

After examining genes likely to decide cell fate early in the differentiation, we next investigated how gene expression changes upon exposure to commonly used astrocyte-specifying growth factors (FGF1/BMP4). We subset our mouse data with both days of AD and the three timepoints of exposure to FGF1/BMP4 (Fig 3A; Fig S13), performing dimensional reduction and clustering as per above (Fig 3B, C). This subset data was analyzed via WOT, identifying cluster 1 as a macrostate (Fig 3D, E) in addition to again recovering a neuronal state (cluster 3; Fig 3F). Cluster 1 expressed a host of genes implicated in astrocyte development, including the RNA-binding protein *Qk*^73^, *Hmgb2*, and the canonical astrocyte marker gene *Slc1a3* (GLAST), in addition to cell cycle genes. This cluster also expressed *Igfbp2*, a molecule implicated in general nervous system development^74^. Cluster 0 expressed *Tgln* and *Tgln2*, genes shown to promote migration and slow proliferation in other developmental systems^75^. These are also two well-known TGF-β target genes^76^, consistent with exposure to BMP4 during the differentiation, which is a major member of the TGFβ signaling family^77^.

**Figure 3.**
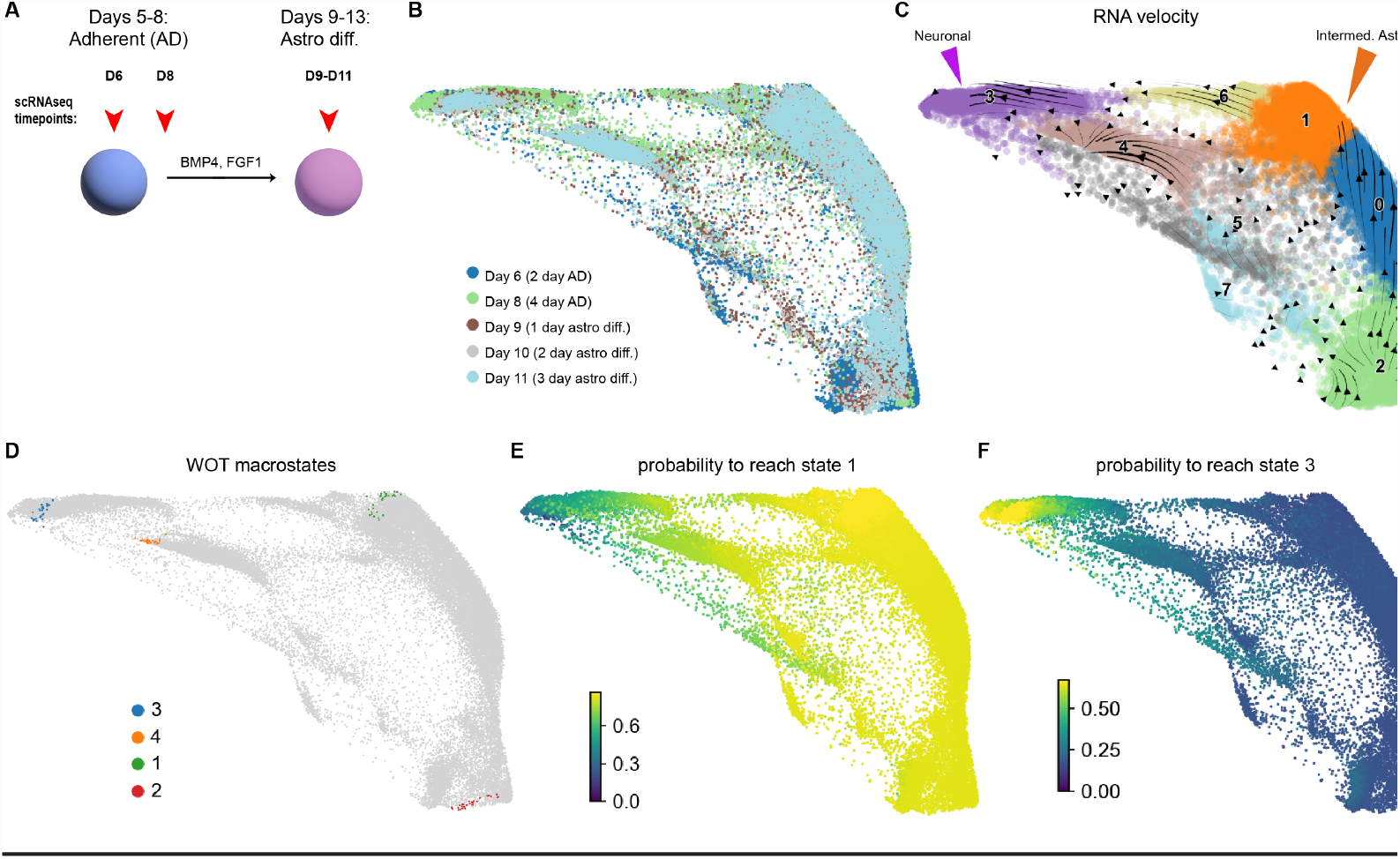
Analysis of subsets of early mouse astrocyte differentiation. **A)**Schematic of mouse data subset and reanalyzed for this figure. **B)** Force-directed graph (FLE) plot of reclustered, subset data colored by batch. **C)** FLE graph with RNA velocity streamlines depicts flow of cell states over time based on splicing ratios for each gene. Neuronal (purple) and intermediate astrocyte (orange) fates are depicted with arrowheads. **D)** FLE graph of the 4 macrostates identified by WOT analysis of this data subset. **E**,**F)** Probability maps for each mature cell type based on the WOT algorithm, scale represents the probability for each cell to eventually enter a given state (state 1 [intermediate astrocyte] or 3 [neuron]).

#### Comparison of potential cell surface marker genes and transcription factor activity

Many groups are interested in cell surface markers as a means to purify glial and neuronal subtypes^36,78^. Accordingly, the early mouse differentiation data was compared with a scRNAseq dataset of early astrocytes acutely purified from 3-day-old mice using the cell surface marker ACSA-2 (*Atp1b2*)^40^. Data from the last five differentiation timepoints (AD and FGF1/BMP4 exposure) were integrated with the 3 replicates of P3 ACSA-2+ astrocytes (Fig 4A,B). RNA velocity analysis of this integrated datasets highlights flow from the putative pre-astrocyte differentiated cell cluster (cluster 0, Fig 4B) towards some of the cells from the P3 dataset (green arrowhead in Fig 4B). Particularly, this cluster (cluster 0) expresses high levels of *Mki67* (Fig 4C), which is consistent with the identification of *Mki67*+ cells as astrocytic precursors in the original P3 dataset^40^. Unlike in the P3 cells, there was minimal expression of *Atp1b2* (ACSA-2) in the differentiated neuronal clusters and intermediate astrocyte clusters (Fig 4D). To investigate alternative candidates for purification of early astrocytes we examined expression of *Slc1a3*/GLAST, the original commercialized astrocyte cell surface marker (ACSA-1; Fig 4E). While heterogenous within P3 cells (Fig 4E; also reported previously in primary mouse brain^79^), *Slc1a3*/GLAST expression was specific for cells that avoided the neuronal fate (red arrowhead in Fig 4B). Finally, we assessed cells for expression of *Gfap*, a traditional astrocyte marker gene (Fig S14D). While we did not observe any *Gfap* expression within our differentiating mouse cells, we did observe heterogeneous *Gfap* expression within the putatively pure P3 astrocyte population (Fig S14D). These observations are consistent with the known variability of *Gfap* expression across astrocytes^80^, and with the lack of *Gfap* expression in the majority of differentiated mouse astrocytes^37^. We next applied BITFAM to these integrated data, and noted cluster-specific TF activity as with the human data (Fig 4F). *Sox9* was active in primary astrocytes (cluster 1), but not in any clusters from differentiated cells; this is consistent with higher *Sox9* gene expression levels detected in primary cells compared to differentiated cells (Fig S14E). Overall, the integrated mouse data reveal potential drawbacks to using the popular ACSA-2 antibody for sorting immature astrocytes as *Atp1b2* (ACSA-2) is induced later during the differentiation compared with *Slc1a3*, in line with the recent identification of *Atp1b2*/ACSA-2 negative glial progenitors in the murine cerebellum^81^.

**Figure 4.**
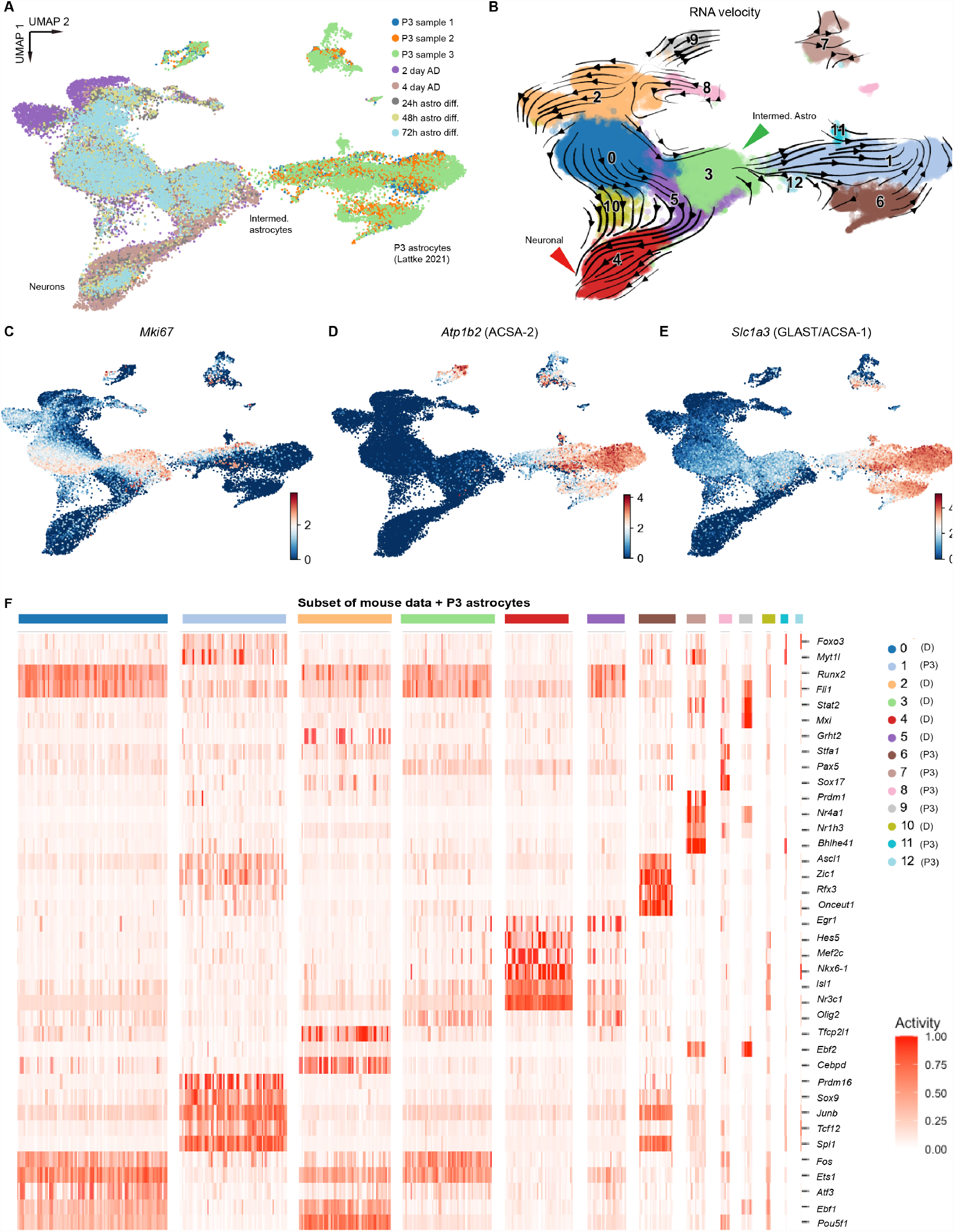
Comparison of putative cell surface marker gene expression patterns in differentiated and primary cells. **A)** Uniform Manifold Approximation and Projection (UMAP) of later mouse differentiation timepoints integrated via Harmony with published postnatal day 3 (P3) mouse astrocyte data. **B)** RNA velocity streamlines are overlain on UMAP with Louvain clusters colored for the integrated data (clustering resolution: 0.6). **C)** Feature plot of *Mki67* expression depicts likely mitotic cells across both datasets. **D)** Feature plots of *Atp1b2*, the gene for the surface protein ACSA-2 used to enrich for astrocytes in the original P3 dataset. Clear heterogeneity of ACSA-2 expression is visible across both datasets. Feature plots for *Sox9* expression, an early astrocyte precursor marker. **E)** Feature plot for *Slc1a3* expression shows medium levels of expression in astrocyte-like differentiating cells, but not in the neuronal-like cells. **F)** Heatmap of BITFAM detected activities for each cluster from panel B (later mouse differentiation timepoints with P3 primary astrocyte data). Clusters are colored on the x-axis corresponding to panel B (legend to the right; D: from differentiated cells, P3: from primary P3 cells), and TFs with detected activity are on the y-axis. P3 data obtained from^40^.

#### Multiomic analysis of two different astrocyte differentiation protocols

We next analyzed two different growth factor combinations used to produce astrocytes from mESCs. To begin, we eliminated the EB stage of the differentiation protocol as there were minimal transcriptional changes during this stage (Fig 2C,D). With this faster, no-EB protocol, we then set up a time series experiment to compare the gene expression changes caused by CNTF exposure versus BMP4/FGF1. Samples were collected every 2 days for 6 days to focus on early changes caused by these growth factors (Fig 5A).

**Figure 5.**
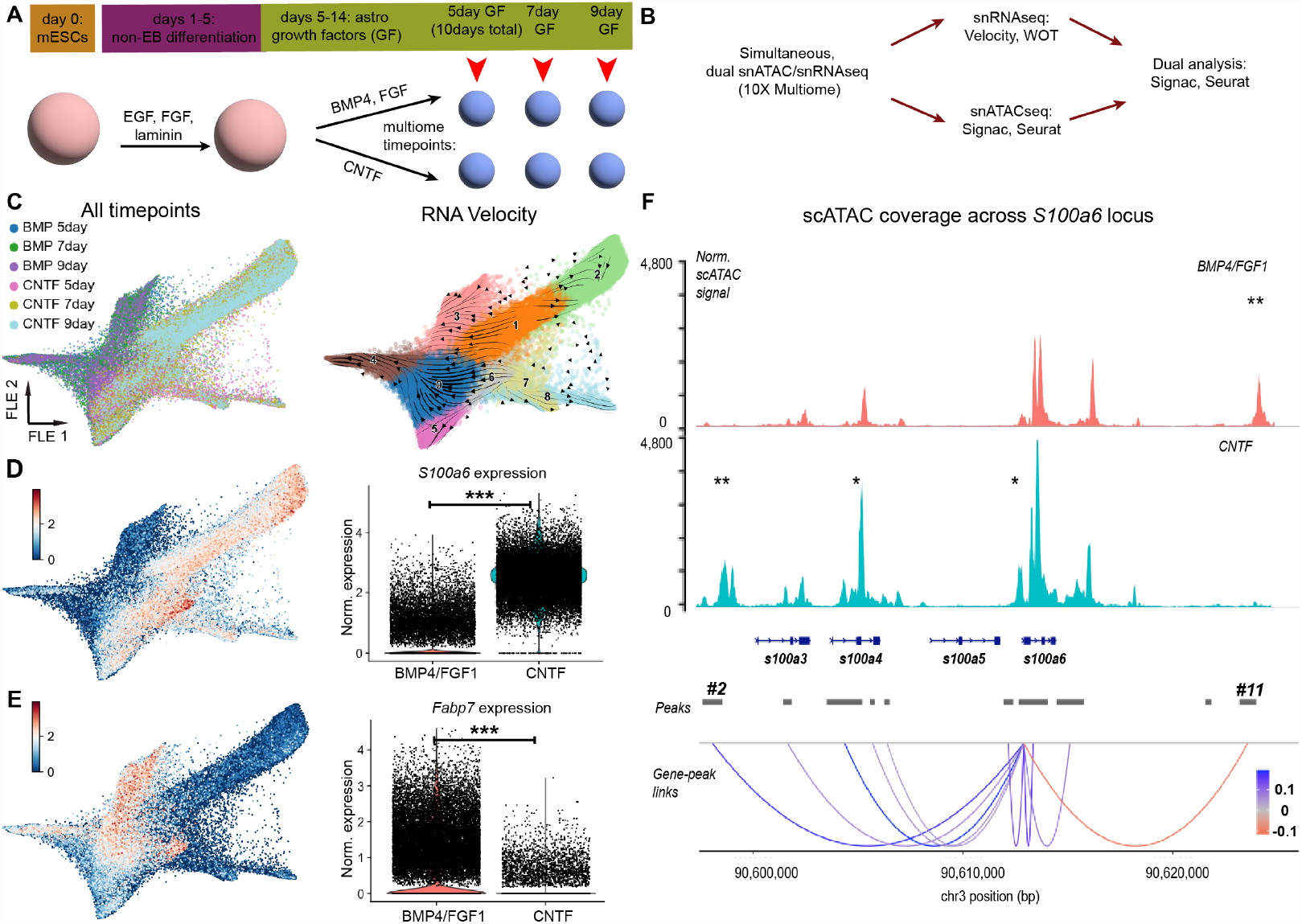
Multiomic analysis of two different astrocyte differentiation time series. **A)** Schematic of modified, no-EB mouse astrocyte differentiation protocol. Time points used for multiomic analysis are marked. **B)** Bioinformatic analysis workflow for multiomic data processing. **C)** (left) Force-directed graph (FLE) of gene expression from 55,137 nuclei collected for multiomic analysis, from 6 time points. See Fig S15 for quality control information. (right) FLE graph with RNA velocity streamlines overlain, to depict flow of cell states over time based on splicing ratios for each gene. Cells were clustered with the Louvain algorithm (resolution: 0.6); clusters are labeled by number on the plot. **D**,**E)** Feature plots and violin plots of marker gene expression for either the BMP4/FGF1 (*S100a6*; D) or CNTF (*Fabp7*; **E)** conditions. ^***^ p < 0.001. **F)** (top panel) Plot of normalized pseudobulk scATAC signal for all cells from each growth factor condition. Peaks are labeled with one asterisk if they have increased signal in one condition versus the other, or two asterisks if the peak is only present in one condition versus the other. (bottom panel) scATAC signal was grouped into peaks using the MACS2 algorithm (see Methods), and peaks were then correlated with gene expression. Correlation scores are highlighted with colored lines in the “Gene-peaks links” panel. Gene body diagrams are from ENSEMBL, as implemented in the “coverage_plot” command in Signac.

To study TF activity and chromatin accessibility changes associated with astrocyte fate acquisition, we processed 55,137 nuclei across 6 time points (Fig 5A; Fig S15) using the 10X Genomics multiomics assay, which allows simultaneous single-nucleus ATAC-seq (snATACseq) and single-nucleus RNAseq (snRNAseq) from the same nucleus (Fig 5B). The data were first analyzed for gene expression and RNA velocity as described above (Fig 5C; Fig S15C,D). Separately, we analyzed gene expression from each differentiation time series on its own (Fig S16), and identified genes correlated with fate acquisition via the WOT algorithm (Fig S16D,E,I,J). We also compared gene expression between the two conditions at the final (6 day) time point, separate from the earlier and thus more immature timepoints (Fig S17).The top marker gene for the CNTF condition at the final, most mature timepoint was *S100a6* (Fig 5D; Fig S17A;C-E), which was recently identified as a marker of a subtype of diencephalic astrocytes in the mouse brain^31^. Overall, this analysis identified differentially expressed genes between two major sets of growth factors used in astrocyte differentiation.

In parallel with gene expression analysis via snRNAseq, we analyzed snATACseq data from the same nuclei. snATAC data were grouped into peaks using the Signac software, then peaks were identified that were enriched in one growth factor condition versus the other. With this list of differentially accessible (DA) peaks, we then applied TF motif analysis to identify TFs specifically active in each growth factor set (Fig S15F,G). Interestingly, in our CNTF condition we noted enrichment of motifs associated with the *Smad* family of TFs (Fig S15G), in line with a recent report implicating *Smad4* in mouse astrogenesis^31^.

To directly connect the gene expression and accessible chromatin analyses, chromatin accessibility was examined at two loci highly correlated with fate acquisition in each growth factor (*S100a6* for CNTF, *Fabp7* for BMP4/FGF1; Fig 5D-F; Fig S15E). Many peaks had significantly different levels of snATAC signal between the conditions (single asterisks in Figs 5F; Fig S15E), and some peaks were fully absent in one condition versus the other (double asterisks in Fig 5F; FigS15E). Using the Signac software package, we calculated a correlation score between accessibility of a peak and expression of the gene of interest. We then plotted the peak-gene links and shaded these links based on the strength of the accessibility-expression correlation. Coverage plots for these genes show significant differences in chromatin signatures at each locus between CNTF and BMP4/FGF1 conditions, mirrored by significant differences in gene expression levels for the two genes (p < 0.01; Fig 5D,E). We next selected a putative DA enhancer peak to delete from the genome and measure whether the absence of the peak altered *S100a6* gene expression (see Methods). We deleted peak #2 (Fig 5F; Fig S15H-J) as it had a high Signac-computed correlation of gene-peak linkage and was located 14,345 bp upstream from the *S100a6* gene body, and thus far away from the promoter. This peak also contained two enhancers previously identified from the ENCODE project based on DNase and H3K27ac scores^82^ (E0736200 and E0736201). Deletion of peak #2 did not significantly affect *S100a6* expression in differentiated cells (Fig S15J). We note that noncoding regions often act in tandem to enhance gene expression, and that deletion of just one peak may not impact gene expression at a detectable level.

Thus, we computationally linked chromatin peaks with gene expression for these genes, and identified areas of open chromatin that may drive glia-specific fate acquisition. Although deleting one of these putative enhancers did not lead to a measurable decrease in gene expression, future studies may be able to test for interactions between multiple noncoding regions that tune gene expression.

#### Integration with mouse developmental cell atlas and functional testing

We next integrated our multiomic gene expression data with a subset of data from a mouse brain development atlas^7^. Samples included: the final two multiomic timepoints (BMP or CNTF day 9), cells from the final timepoint of the differentiation with EBs, and cells from Lattke et. al. 2021 (7,042 cells/nuclei total), together with 109,700 cells from the mouse cell atlas spanning e15.5 to e17.5 (Fig 6A-C). After clustering and integration via Harmony, at low resolution the differentiated cells and nuclei clustered together with groups of cells from the primary mouse atlas cells (dashed boxes; Fig 6B,C). Higher resolution clustering uncovered marker genes that separated the primary cells from the differentiated cells, although many marker genes were still shared between differentiated and primary cells at higher resolution (e.g. *Sparc*; Fig S18B-G). Differentiated cells mainly clustered with glioblasts and pineal gland cells from the primary atlas (Fig S18E), suggesting that the new differentiation protocol produces cells that are transcriptionally similar to early (e15.5-e17.5) primary mouse astrocytes.

**Figure 6.**
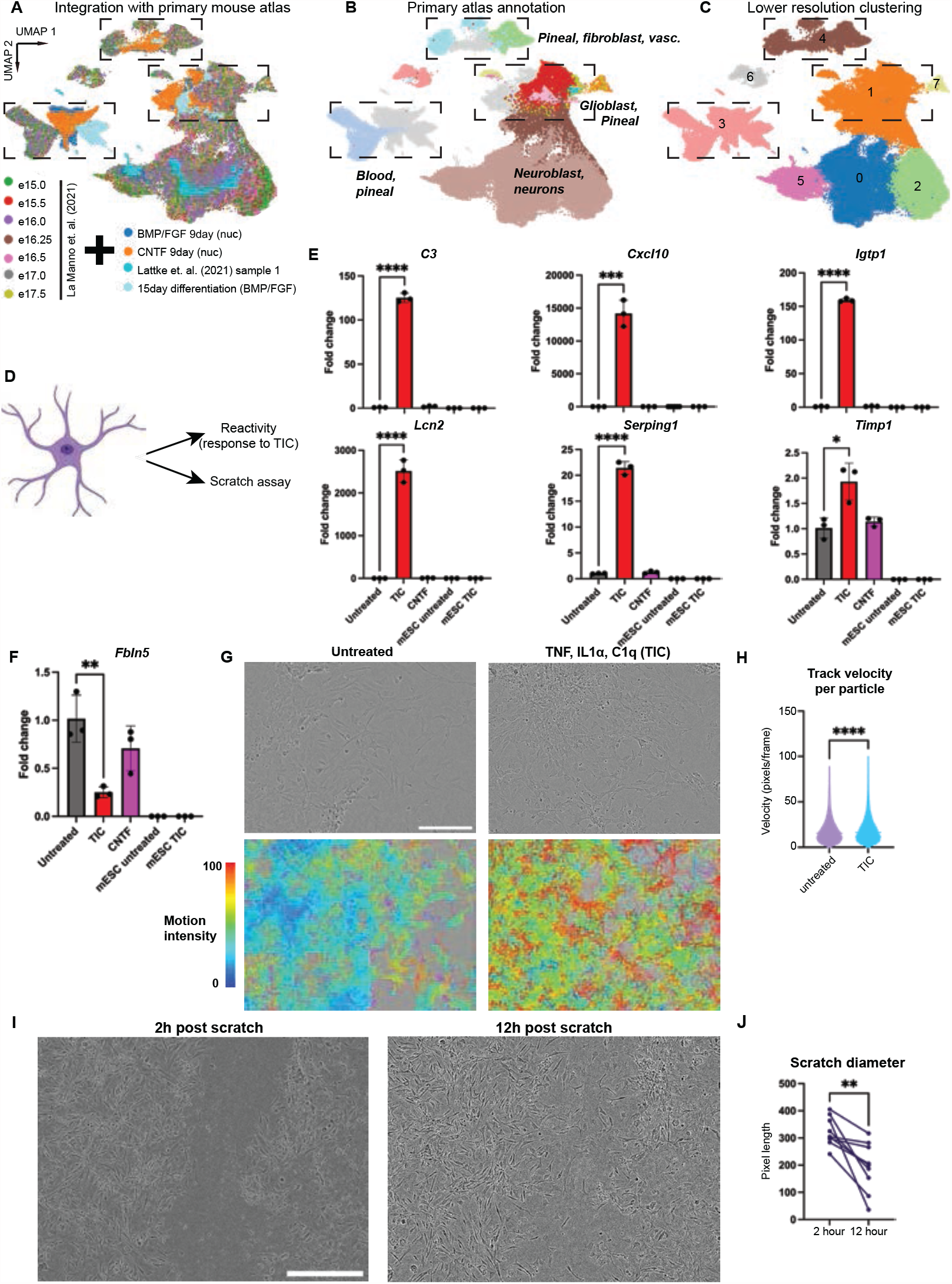
Data integration and functional analyses of differentiated immature astrocytes. **A)** UMAP plot of final mouse differentiation timepoints integrated with mouse embryonic brain dataset^7^. Samples were integrated using Harmony to correct for different scRNAseq technologies; 109,700 cells remained after quality control filtering. Dashed boxes on the UMAP plot highlight areas of overlap between differentiated cells and cells from the primary atlas. **B)**Atlas cells from A) labeled with annotations from the primary cell atlas. Select groups are highlighted; full annotation plot can be found in Fig S18A. **C)** Cells were clustered using the Louvain algorithm at low resolution (0.1), and differentiated cells cluster with a subset of embryonic mouse brain cells; marker genes for the dashed clusters are presented in Fig S18D, with feature plots for select genes in Fig S18E,F. D) Overview of astrocyte functional assays tested. **E)** Quantitative Reverse Transcription PCR (qRT-PCR) results of differentiated immature astrocytes exposed to TIC (TNF, IL1a, and C1q,) or CNTF (control) demonstrates dramatic induction after 24 h of many genes previously identified as upregulated in primary mouse brain after lipopolysaccharide (LPS) injection^2^. T-test between no TIC and TIC treated samples, n=3 independent culture wells, asterisk represents p < 0.01 for all samples. **F)** qRT-PCR data for one gene with decreased expression in TIC-treated differentiated astrocytes after 24h, *Fbln5* (statistical analysis as in previous panel). **G)** Single frames from chronic imaging of TIC-treated cells (top) with CellTracker motility analysis overlay (bottom). **H)** Analysis of chronic imaging via CellTracker shows increased motility in TIC-treated cells compared to untreated controls (p < 0.0001, statistical analysis as in panel E). **I)** Single frames from chronic imaging of cells subjected to the scratch assay 2 (left) and 14 (right) h after scratch. Contrast and brightness identically enhanced in both frames for display purposes. **J)** Quantification of scratch repair over time demonstrates significant decreases in scatch diameter over 12 h (p < 0.01, statistical analysis as in panel E). Quantification example can be found in Fig S18I.

Given the transcriptional similarity between the differentiated cells and primary mouse cells, the functional properties of differentiated cells were tested using two in vitro assays (Fig 6D). To begin, we examined whether the differentiated astrocytes could respond to an inflammatory cocktail that induces reactivity in astrocytes^83^. After 24 hours of exposure to TNF, IL1a, and C1q (TIC), the differentiated cells dramatically and significantly upregulated reactive astrocyte marker genes *Cxcl10, C3*, and *Serping1* (Fig 6E). In contrast to previous work from our group, the differentiated cells down-regulated *Fbln5* after 24 h of stimulation, unlike primary astrocytes collected 24 h after systemic injection of the bacterial cell wall endotoxin lipopolysaccharide (Fig 6F). This gene is thought to promote cell-cell adhesion^84^, so given its downregulation we measured cell motility in response to TIC. Chronic imaging of TIC-treated cells compared to PBS-treated controls showed an increase in cell motility and cellular process motility, which we quantified using an automated image tracking software (Fig 6G,H; see Methods). We further assessed cell motility through a scratch assay using the same chronic imaging methods (Fig 6I,J; Fig S15E,F; see Methods). The mESC differentiated astrocytes are able to rapidly migrate and extend processes across the scratch wound (Fig 6I,J). Thus, our differentiated astrocytes display at least two functions of mature astrocytes: changes in gene expression and motility in response to reactive factors, and successful repopulation following a scratch wound.

## DISCUSSION

Here, we report complementary bioinformatic analyses of sc/snRNAseq data to parse the temporal order of gene expression during glial development in both mouse and human glial differentiation protocols. We provide a set of hypotheses about gene expression in progenitor cells and the order of events during differentiation to test in future lineage tracing experiments, both in vitro and in vivo. Bioinformatic analysis of a rich scRNAseq dataset (∼298,000 cell/nuclei transcriptomes newly sequenced, and ∼120,000 published cells from other groups) enabled us to identify likely glial-lineage-committed precursors, and generally to explore the genes involved in specification of neurodevelopmental fate in mice and human.

In our human scRNAseq dataset, we identified final, stable macrostates and calculated probabilities for each intermediate cell to reach one of the macrostates. This approach revealed that astrocytes and neurons likely share common progenitor cells in this differentiation model, and that oligodendrocytes are produced after astrocytes; both observations are consistent with the progression of development in vivo. Additionally, we were able to identify genes highly correlated with high probabilities for cells to enter a given macrostate over the others. We suggest that *TTYH1* is a strong candidate for a driver of astrocyte fate specification, given its high correlation with astrocyte probability and its purported ability to signal through the Notch pathway^57^, thought to be important for the acquisition of astrocyte identity^25^. Past work has localized *TTYH1* to cytoskeletal processes in glioma cells, and shown that its presence leads to tighter networks of tumor cells58; perhaps the same phenomenon occurs normally during astrocyte development.

In our mouse differentiation data, we found an early “off-ramp” fate for a subset of cells that will become neurons, consistent with past reports of transient progression through neuronal fate during astrocyte differentiation^37^. We observed heterogenous expression of popular astrocyte sorting markers during differentiation, suggesting caution when designing experiments using these markers. In many cases, gene expression was similar in both the human and mouse differentiations, including early expression of *Dcx*/*DCX* followed by shared expression of putative astrocyte precursor genes like *Hmgb2*/*HMGB2* and *Slc1a3*/*SLC1A3* across both species. There was some species divergence, for example with *Nhlh1*/*NHLH1*, which is suggested to define neuron-committed progenitors in mouse but was expressed in human astrocyte progenitors, and also divergence in the absence of *Gfap* expression during early mouse differentiation. This is a major difference between in vivo and in vitro progression of astrocyte differentiation first noted by others^37^ and replicated in our dataset. Overall, the lack of early *Gfap* expression in differentiation is a striking difference between in vivo and in vitro neural development, given the historical emphasis on the study of *Gfap*+ radial glia as the earliest mammalian neural progenitors^13^. In order to better compare in vitro versus in vivo glial fate acquisition, in Fig 6 we 1) integrated our data with a mouse developmental brain atlas, and 2) tested select astrocyte functions in cell culture assays to show that our differentiated immature astrocytes partially recapitulate primary cells. Similar functions have already been demonstrated for the human differentiation protocol in past work^36,78^.

Although the measurement of gene expression is clearly a useful tool for analyzing the progression of glial differentiation, TFs are ultimately the key determinants of cell fate choices^85,86^. Thus, we applied the BITFAM algorithm to our datasets and examined mechanisms underlying gene expression differences during glial development via multiomic analysis, where we identified regulatory sequences correlated with glia fate selection. It will be important to compare these genomic areas involved in astrocyte differentiation with those areas identified in states of astrocyte reactivity and disease, as a baseline to uncover specific mechanisms driving health-relevant transcriptional changes in astrocyte states. This dataset, which to our knowledge is the first single-cell/single-nuclei time series analysis of directed glial differentiation in either rodents or humans, will provide a valuable resource for researchers interested in studying these processes. The genes identified as key drivers of specific lineages may also guide groups interested in further optimizing differentiations, either to improve purity of differentiation protocols (e.g., minimizing the production of cells from non-desired lineages) or to differentiate specific subtypes of astrocytes for functional testing, a burgeoning area of interest. We also characterized areas of the genome that may be critical for regulation of astrocyte fate acquisition, in addition to genomic regions that are associated with expression of reactive genes like those seen in CNTF treatment. Disruption of these genomic areas might block the expression of reactive genes in astrocytes, a topic of enormous interest and importance.

## METHODS

### Cell culture and differentiation

#### Human iPSC culture

Human iPSCs for all experiments, except the one displayed in Fig S2, were from a single line originally reprogrammed at NYSCF from skin fibroblasts of a healthy female individual (age 52 at collection), using the fully automated NYSCF Global Stem Cell Array® platform^42^. Following our published protocol, we maintained human iPSCs in mTeSR1 medium in a 37 °C incubator with 5% CO2^88^s. To prepare for induction, we performed enzymatic digestion using Accutase and plated 1.5×105 iPSCs per well on Geltrex-coated 6-well plates, in maintenance medium with 10 μM Y27632 for 24 hours. For astrocyte differentiation, we used serum-free medium with patterning agents as previously described^36,88^ and schematized in Fig S1A. At day 20, we plated 50 neu-rospheres per well into poly-ornithine/laminin-coated 6-well plates. Spheres attached to the matrices within a few hours and neural progenitors migrated out and spread across the surface area of the well. At days 30 and 50, we collected single cells using papain (Worthington LS003126) digestion and prepared cells for scRNAseq analysis. For the final time point corresponding to mature astrocytes, we used transcriptomic data from the same iPSC line generated in our previous study^88^.

For our multi-line experiment, we used 9 iPSC lines from an iPSC collection recently generated at NYSCF to enable large scale studies on Alzheimer’s disease^89^. Peripheral blood mononuclear cells from individuals enrolled in the Religious Order Study (ROS) or the Rush Memory and Aging Project (MAP) were reprogrammed using Sendai virus. iPSC line generation and quality controls were performed using the NYSCF fully automated Global Stem Cell Array technology. iPSCs from each line were differentiated in parallel following the same protocol as above. At day 73, we collected nuclei for processing with 10X multiome as follows: Cells were dissociated with TrypLE (Gibco 12-605-010) for 15 min at room temperature, then resuspended in 1 mL EZ lysis buffer (Sigma (NUC101-1KT)). Cells were triturated ten times with a P1000, then left on ice for 10 min. EZ buffer was washed out with 3 washes in PBS + 0.5% BSA, then filtered through a Flowmi 40 µm pipette tip filter (VWR H13680-0040) before counting.

#### Mouse ESC culture and differentiation

Mouse ESC experiments were performed on two separate mESC lines previously validated elsewhere: A17iCre^90^ and BL6/CAST Δ*Piga*^91^. Cells from both lines were maintained in 80/20 media for expansion and freezing, in a 37 °C incubator with 5% CO2. This media contains 80% 2i mESC media^69^, and 20% general mESC media. Specifically, this 80/20 media contains 40% Advanced DMEM/F12 (Gibco 12-634-010), 40% Neurobasal (Gibco 21103049), 10% Knockout DMEM (Gibco 10829018), in addition to 2.5% fetal bovine serum (Corning 35-075-CV), 0.75x N2 nutrient mix (Thermo 17502048), 0.75x B27 nutrient mix (Thermo 17504044), 0.75x L-Glutamine (2mM; Gibco 25-030-081), 1X beta-mercaptoethanol (Gibco 21985023), 1x non-essential amino acids (Gibco 11140050), 1x nucleosides (Millipore ES-008-D). This media also contained leukemia inhibitory factor (LIF) at 1000 units/mL (Millipore ESG1107), 3 µM CHIR99201 (Biovision 1991-1), and 1 µM PD0325901 (Sigma PZ0162-5MG) to maintain pluripotency during the mESC stage.

Mouse differentiation was performed with a protocol derived from^38^, and is schematized in Fig 2A; Fig S9A. On day 1 of the main differentiation (with EBs), mESCs were trypsinized with TrypLE Express (Gibco 12-605-010) for 2 min, quenched with 80/20 media, then replated in low-adherence 10 cm tissue culture dishes (Corning 3262) at densities of 1.5 million cells per dish in AK media (50% Neurobasal (Gibco 21103049), 50% Advanced DMEM/F12 (Gibco 12-634-010), 1X L-Glutamine (Gibco 25-030-081), 13% Knockout Serum Replacement (Gibco 10-828-028)). At days 3 and 5 of the differentiation, embryoid bodies were collected in a 50 ml tube and allowed to settle via gravity for 20 min. AK media was refreshed and cells were replated back into the same low-adherence 10 cm dish. At day 6, embryoid bodies were collected via gravity, then trypsinized in 4 mL TrypLE for 5 min with intermittent swirling by hand. TrypLE was quenched with AK media, and cells were filtered through a pre-wet 70 µM nylon cell filter. Filtered cells were replated on poly-d-lysine (50 µg/mL; Sigma, P6407)- and laminin (10 µg/mL; Sigma, 11243217001)coated 10 cm tissue-culture dishes at 1 million cells/dish, and murine EGF (Peprotech 315-09, 20 ng/ mL), murine FGF1 (Peprotech 450-33, 10 ng/mL), and laminin (Sigma 11243217001, 1 µg/mL) were added to AK media when plating cells for the adherent differentiation phase. Media was replaced every other day until day 11, when cells were switched to differentiation media, AK with FGF1 (Peprotech 450-33, 10 µg/mL) and BMP4 (Peprotech 315-27, 10 µg/mL). Cells were replated once more onto PDL+laminin dishes upon reaching 70% confluency (∼day 15), and maintained in differentiation media (AK plus FGF1, BMP4) replaced every other day. CNTF (Peprotech 450-50, 10 µg/ mL) was used as an alternate to FGF1/BMP4 where indicated.

For the non-EB differentiations, the differentiation was as follows. For the adherent differentiation (AD) stage, cells were plated (15,000 cells/well) on gelatin- coated 6-well TC dishes in AK media + EGF (Peprotech 315-09, 20 ng/mL), FGF1 (Peprotech 450-33, 10 ng/mL), and laminin (Sigma 11243217001, 1 µg/mL). Media was refreshed the day after plating, and then replaced every other day following. After 5 days total, media was changed to astrocyte growth factor media, AK + FGF1 (Peprotech 450-33, 10 µg/mL) and BMP4 (Peprotech 315-27, 10 µg/mL), with replacement every other day for another 6 days until harvesting for 10X scRNAseq. Cells were replated at 100,000 cells/well on gelatin-coated 6-well dishes with growth factors and laminin once they reached 80% confluency.

#### Engineering Δpeak #2 mESC line

We designed guide RNAs targeting PAM sites for Cas9 bordering the left and right sides of peak 2 (chr3:90597589-90598549, mm10) from Fig 5F. A repair ssODN was designed with homology arms to each of the cut sites. We cloned these guide RNAs into vectors expressing SpCas9 and either a puromycinor a blasticidinresistance marker gene, and applied dual selection to 1.5×10^6^ cells 24 h after transfection using the Lonza 2b nucleofector. Selection was performed for 48 h total, then we isolated individual surviving colonies for genotyping using previously published methods^91^. Genotyping was confirmed by Sanger sequencing.

#### Reactive stimulus imaging and qRT-PCR

Astrocytes from the final 9-day differentiation timepoint BMP/FGF4 differentiation were stimulated using a cytokine cocktail previously shown to induce reactivity in astrocytes^83^. After addition of TIC (TNF, IL1a, and C1q,) cultures were imaged every 30 min at 20x for 24 h using an Incucyte S3 incubator microscope. Still images were exported and analyzed in ImageJ, with quantification schematic outlined in Fig 6F. Thresholds were applied across all images in ImageJ v2.1.0 and movement was analyzed with the Trackmatev6.0.3 plugin^92^. A Laplacian of Gaussian filter was applied to recognize spots with a 15-pixel blob diameter. Spots were then tracked through frames using a Linear Assignment Problem tracker and maximal velocities per spot were extracted.

#### Quantitative Real-time PCR (qRT-PCR)

RNA was extracted using the RNEasy Plus kit (Qiagen) with gDNA removal columns, reverse transcription was performed on 100 ng of RNA using the Qscript mastermix (Quanta Biosci 101414-108), and qRT-PCR was performed on a Roche 480 Lightcycler with KAPA SYBR FAST (Kapa Biosystems KK4610) after plate loading with an Echo 550 liquid handler. qRT-PCR was data was collected from 3 wells from at least 2 separate differentiations, and run in technical triplicates in 384 well plates. Expression was calculated using the ΔCt method; Ct values for each gene were subtracted from the housekeeping gene (Gapdh) Ct value for that sample. ΔCt values for each condition were averaged, and experimental (TIC or CNTF treated) values were subtracted from control (untreated) to produce ΔΔCt. This final value was converted to fold change relative to untreated by taking 2 raised to the - ΔΔCt.

All qRT-PCR primers were designed to span exon-exon junctions, and no-RT and water controls were run for all reactions to ensure full removal of gDNA. Primers used were as follows: *S100a6*: F CGACACATTCCATCCCCTCG, R CACTGGGCTAGAAGAAGCGCA; *Fbln5*: F CTTCAGATGCAAGCAACAA, R AGGCAGTGTCAGAGGCCTTA; *C3*: F TCACTATGGGACCAGCTTCA, R TGGGAGTAATGATG- GAATACATGGG; *Cxcl10*: F CCCACGTGTTGAGATCATTG, R CACTGGGTAAAGGGGAGTGA; *Timp1*: F AGTGATTTC- CCCGCCAACTC, R GGGGCCATCATGGTATCTGC; *Serping1*: F ACAGCCCCCTCTGAATTCTT, R GGATGCTCTC- CAAGTTGCTC; *Igtp1*: F GTAAGGCTTCTGAGCAGGTTCT, R TATGGAGTATGAAGGTCTATGTCTG; *Lcn2*: F CCAGTTCGCCATGGTATTTT, R CACACTCACCACCCATTCAG; *Gapdh*: F AGAACATCATCCCTGCATCC, R CACATTGGGGGTAGGAACAC.

#### Scratch assay

A monolayer of astrocytes from the final 9-day rapid differentiation timepoint were scratched using a P200 pipette tip. Cultures were imaged every 30 min for 24 h using an Incucyte S3 incubator microscope. Still images were exported and analyzed in ImageJ, with quantification schematic outlined in Fig S18E. For Fig 6J, scratch distance was quantified for three separate wells scratched concurrently, with 3 areas measured per scratch; a t-test was run in Prism 9 (GraphPad Inc.) to test for significant changes.

### Single cell and nuclei RNAseq and ATACseq

#### 10X scRNAseq data collection & sequencing

For collection of 10X scRNAseq samples, media was re-placed 3 h prior to 10X processing for all cultures. Mouse cells were trypsinized for 10 min with TrypLE, then quenched with appropriate media. Cells were triturated with a 10 mL serological pipette, filtered with a 40 µM filter, and diluted to 1000-1200 cells/µl. Human samples were dissociated according to a modified papain-based protocol (Worthington), then filtered through a 40 µM filter before counting and dilution for scRNAseq.

17,400 cells were loaded onto 10X chip G (mouse samples) or chip B (human samples), and run with v3.1 3’ chemistry (mouse samples) or v3.0 3’ chemistry (human samples). Library preparation was performed according to manufacturer’s instructions, and sequenced using a NOVASeq6000 sequencer (Illumina).

#### 10X multiomic data collection & sequencing

For collection of 10X multiomic samples, media was replaced 3 h prior to nuclei collection. Mouse cells were trypsinized for 10 min with TrypLE, quenched, and spun as for scRNAseq collection. The pellet was washed with 5 mL PBS once, then spun again. The washed pellet was then resuspended in 500 µL ice-cold Sigma EZ Nuclei Extraction Buffer (Sigma-Aldrich), pipetted 10X with a P1000 pipette, then left on ice for 10 min. Nuclei were pelleted at 500 rcf for 5’, washed once with 500 µL PBS+0.5%BSA, then resuspended in 10X Genomics Nuclei Buffer. 17,400 nuclei per sample were used as input for scATAC, snRNAseq and ensuing library prep, which were performed according to manufacturer’s instructions and then sequenced on a NOVASeq6000 sequencer (Illumina).

### Bioinformatic analysis

#### 10X scRNAseq data processing and quality control

All analyses were executed locally on a laptop computer, except for Cell Ranger, which was performed on NYU Grossman School of Medicine’s Ultra Violet Cray supercomputer cluster. Fastq files from sequencing were demultiplexed, then used as input for Cell Ranger (v6.0.1) for assignment of reads to single cells. Filtered count matrices from Cell Ranger were then used as input to the Muscat quality control pipeline in R^43^. After doublet detection using scds (v1.1.2) and calculation of the mitochondrial gene contribution using scater (v1.13.23) within the SingleCellExperiment library (v1.7.11), cells were filtered using the median absolute deviation (MAD) module where cells containing a greater than 2.5 MAD in feature counts, number of expressed features and percentage of mitochondrial genes were discarded. Between 65 and 85 percent of cells were kept post-filtering, see Figs S1B, S7E for details and numbers for each sample.

Barcodes of cells that passed Muscat quality control were used to filter the .loom file produced by the velocyto python package^46^, which analyzes the .bam file output of CellRanger for splicing to prepare for RNA velocity analysis. This filtered .loom file was used as input to the Scanpy^87^ pipeline, implemented as part of the scvelo pipeline^47^. All samples were normalized and filtered according to standard Scanpy parameters: genes were filtered to only retain genes with number of spliced counts >20, and then the most variable remaining genes were identified by the pp.filter_and_normalize command. For our human data, we kept the top 2000 most variable genes for further analysis; for mouse, we kept the top 4000 genes. We performed principal component analysis (PCA) on the most variable genes via the pp.pca command, then data were integrated along the ‘batch’ variable via implementation of the Harmony algorithm in the Scanpy external API (pp.harmony_integrate); neighbors were found via pp.neighbors using 40 PCs, 40 neighbors, and using X_pca_harmony; cells were clustered using the Louvain algorithm implemented in scvelo (scv.tl.louvain) at resolution values ranging from 0.3 to 1.6, then dimensionality reduction was performed via scv.tl.umap or using a force-directed graph (sc.tl.draw_graph); and finally, marker genes (i.e., top-positively-enriched genes) for each Louvain cluster were found using sc.tl.rank_genes_groups with a Wilcoxon ranksum algorithm.

For pseudobulk analysis, python objects were converted into Seurat objects using the SeuratDisk package (v0.0.0.901). Samples were then split into individual timepoints using the Seurat (v4.1.0) SplitObject() command. Pseudobulk data was calculated for pre-normalized expression data for each of the overall most variable genes using the AverageExpression() command, as recommended in Seurat documentation.

#### RNA velocity analysis

The .loom file from velocyto was used to compute RNA velocities for each cell according to standard parameters for the software. ScVelo produces both stochastic and dynamic models of RNA velocity, which we compared by computing a consistency score for each cell, for each modeling method, as recommended by the authors. Pseudotime was then computed based on RNA velocity results, and latent time was inferred via dynamic velocity results.

#### Waddington Optimal Transport (WOT) analysis

The WOT algorithm^11^ was used via its implementation in CellRank. Batch names were changed to sequential numbers, and then a WOT kernel was initialized via the WOTKernel command. Growth rates were calculated by compute_initial_growth_rates, which uses the scanpy growth score (a gene module score for cell cycle genes). A transition matrix was calculated partially using these growth rates as a basis via compute_transition_matrix with 3 growth iterations. We found that the transition matrix was nearly identical whether or not we included growth rates. We then computed a connectivity kernel using the ConnectivityKernel command, which calculates a diffusion pseudotime score for each cell. These kernels were combined at weight of 0.9 for WOT, 0.1 for the connectivity kernel, as recommended by the original authors. We saw minimal differences between our outcomes whether or not we included the 0.1 weighted connectivity kernel. We then used a Generalized Perron Cluster-Cluster Analysis (GPCCA) estimator to define macrostates, i.e. cell states with metastability. We then used a Schur decomposi-tion to compute a coarse transition matrix that enabled us to compare the stability and likely temporal order of the various macrostates. We selected macrostates for further analysis based on prior knowledge of expected cell types (humans: oligodendrocyte, astrocyte, neuron; mouse: astrocyte-precursor, neuron), and then produced absorption probability plots/fate maps for these macrostates with the plot_absorption_probabilities command. We took the logarithm of these probabilities to compute log-odds for each macrostate, and compare across states for the log-odds plots; in these plots, we colored cells red based on normalized expression of a gene of interest as explained in the text/figure legend. Finally, we computed genes correlated with high probability to become a certain macrostate via the compute_lineage_drivers command. This correlation is a Fisher transformation of the Pearson’s coefficient between gene expression and macro- state probability, as discussed in the CellRank manuscript.

#### BITFAM algorithm

We exported the count matrices from our Scanpy objects for use with the BITFAM algorithm (v1.0) implemented in R, using standard settings. BITFAM results were first visualized as a t-SNE using the Rtsne R package (v0.15). Finally, cluster IDs were exported from Scanpy and used to create a heatmap of inferred TF activity for each cluster, following sample code from the BITFAM package.

#### Analysis of Lattke et. al. 2021 and La Manno et. al. 2021 published scRNAseq data

Raw .fastq files were downloaded from P3 ACSA-2 purified astrocytes published in^40^, NCBI GEO #GSE152223. The fastq files for the raw data were analyzed via CellRanger v6.0.1, and then count matrices were imported into muscat for processing (see above). We used the published raw data to produce .loom files via Velocyto splice calling, which was not done in the original analyses. We then processed these.loom files for RNA velocity as above, filtered for quality control with Muscat, and then we combined both published datasets with our later mouse timepoints via the Harmony algorithm. For integration with the primary mouse brain atlas^7^, we downloaded .loom files from the Linnarsson lab website and then used Harmony to integrate with our datasets.

#### Multiomic analysis

Multiomic data was preprocessed by CellRanger ARC software (v4.0.5), then initially the scATAC and snRNAseq datasets were processed separately, in Seurat^93^/Signac^94^ and ScVelo/CellRank respectively. For scATAC, nuclei QC was based on thresholds on the following QC metrics: nCount_ATAC < 120000, nCount_ATAC > 500, nCount_RNA < 50000, nCount_RNA > 500, TSS.enrichment > 1 (Fig S15A). scATAC signal was transformed into peaks by the MACS2 peaks calling algorithm, as suggested by Signac authors, then added to the Seurat object as an assay. Peaks were analyzed for transcription factor motif enrichment (Fig S15F,G) after finding differentially accessible peaks with logistic regression (FindMarkers with ‘LR’ option for test.use), and based on analysis with the JASPAR2020 dataset. Peaks were also linked to gene expression using the Signac software, which generates a z-score for each peak within 500kB of a gene’s TSS compared to a background set of correlation coefficients for 200 peaks randomly selected from a different chromosome. Gene coverage plots for Fig 5F and Fig S15E were generated using the CoveragePlot() command, which plots pseudobulk scATAC coverage.

#### Immunohistochemistry

Immunofluorescence analysis was performed at the end of the human differentiation to assess the presence of astrocytes (GFAP), oligodendrocyte lineage cells (SOX10) and neurons (MAP2) within the cultures from all 9 ROS/MAP lines, as previously described^78^. Briefly, cells were fixed in 4% paraformaldehyde for 10 min and then washed 3 times in DPBS. Next, cells were incubated for 1 h in blocking solution (5% donkey serum, 0.1% Triton-X-100 in DPBS) and overnight with the following primary antibodies: goat SOX10, 1:100 (R&D system, #AF2864); chicken MAP2, 1:1000 (Abcam #Ab5392; mouse GFAP, 1:1000 (EMD Millipore #MAB360). The morning after, cells were incubated 1 h in secondary antibodies (Anti-goat 555, Invitrogen #A21432, anti-chicken 647, Jackson ImmunoResearch #703605155, anti-mouse 488, Invitrogen #A32766), washed 3x in DPBS and then incubated with Hoescht 33342 (1:1000 dilution) for 10 mins. Images were acquired using a Zeiss scanning confocal microscope (LSM800).

Mouse cultures were fixed in 4% paraformaldehyde (Electron Microscopy Services) for 15 min at room temperature (RT), then plates were washed 3x with phosphate-buffered saline (PBS) before blocking in PBS + 0.1% Triton-X and 3% goat serum for 1 h at RT. Primary monoclonal antibodies to GFAP (ThermoFisher #13-0300, 1:500) or S100A6 (Abcam #ab181975, 1:2000) were applied in blocking solution overnight at 4 oC with rotation. One well per plate was processed without primary antibody as a negative control. The next day, plates were washed quickly 3x with PBS, then washed 3x more with 10 min rotation. Secondary antibodies (Invitrogen, 1:2000) with 4′,6-diamidino-2-phenylindole (DAPI; ThermoFisher, 1:5000) were applied in blocking solution for 1h at RT on an orbital shaker. Plates were washed again as with the primary antibody, then imaged using a Keyence BZ-X700 fluorescent microscope. Channels were separated to make figures using Fiji/ImageJ software.

### Statistics

Clustering was calculated by the Louvain algorithm. Marker genes for each cluster were calculated using the Wilcoxon rank-sum algorithm implemented in Scanpy. Statistical tests used in the WOT and BITFAM algorithms are discussed in those sections of methods. Student’s t-tests for Fig 6 were run in Prism 9 (GraphPad). Sample sizes were determined based on previous publications, and independent biological replicates range from 1 to 3 for all experimental modalities used in this study. All statistical tests used are highlighted in the legend of each figure. No data were excluded from the analyses, except when performing quality control filtering as discussed above. The experiments were not randomized. The investigators were not blinded to allocation during experiments and outcome assessment.

### Reproducibility

For human data, we separately loaded single-cell suspensions from two adjacent wells of a 6-well plate in individual 10X lanes, processed each lane as a separate library, then combined the lanes for analysis bioinformatically after checking for any batch effect; the 30-day and 50-day timepoints were from two separate cultures differentiated on separate dates. For mouse data, EB timepoints were collected from two independent cultures first, to check for replicability/batch effects (Fig S1E). Once convinced of minimal batch effects, we then collected all remaining time points on the same day and processed each condition in its own 10X scRNAseq lane. Accordingly, each time point is from an independent differentiation culture (i.e., each timepoint began from frozen mESC vials on a different day, staggered for collection on the same day). Further, we compared immunostaining between two mouse stem cell lines under the same differentiation conditions (Fig S7G). We performed all of our human scRNAseq experiments in one cell line for Fig 1 and associated supplements (except the multi-line comparison in Fig S2), and performed mouse scRNAseq experiments in two different cell lines as detailed above.

## Acknowledgements

The computational requirements for this work were supported in part by the NYU Langone High Performance Computing (HPC) Core’s resources and personnel. Funding for this work was provided by NIH/NEI (R01EY033353), the Cure Alzheimer’s Fund, MD Anderson Neurodegeneration Consortium, Anonymous Donors, the Blas Frangione Foundation, and The Alzheimer’s Association (SAL); we also acknowledge the generous support of Paul Slavick and the Parekh Center for Interdisciplinary Neurology (SAL); the Dark Matter Project funded by an NIH Center for Excellence in Genome Science – Grant IDs 5RM1HG009491, 3RM1HG009491-03S1, 3RM-1HG009491-03S2 (JDB); National Institute of Neurological Disorders and Stroke (NINDS) grant 1R21NS111186 (VF); NINDS T32 (NYU/Dasen) 5T32NS086750 (PWF).

## Competing Interests

SAL is an academic founder and sits on the SAB of AstronauTx Ltd., and a SAB member of the BioAccess Fund. JDB is a Founder and Director of CDI Labs Inc., a Founder of and consultant to Neochromosome Inc., a Founder, SAB member, and consultant to ReOpen Diagnostics LLC, and serves or served on the SAB of the following: Sangamo Inc., Modern Meadow Inc., Rome Therapeutics Inc., Sample6 Inc., Tessera Therapeutics Inc., and the Wyss Institute.

## Notes

### Summary of Updates

Substantial changes were made during peer review, including new functional data and published dataset integrations in fig 6. Some revision experiments were performed in collaboration with co-authors added during peer review.

