## Supplemental Figures 1-18 for "Time series scRNAseq analysis in mouse and human informs optimization of rapid astrocyte differentiation protocols"

A

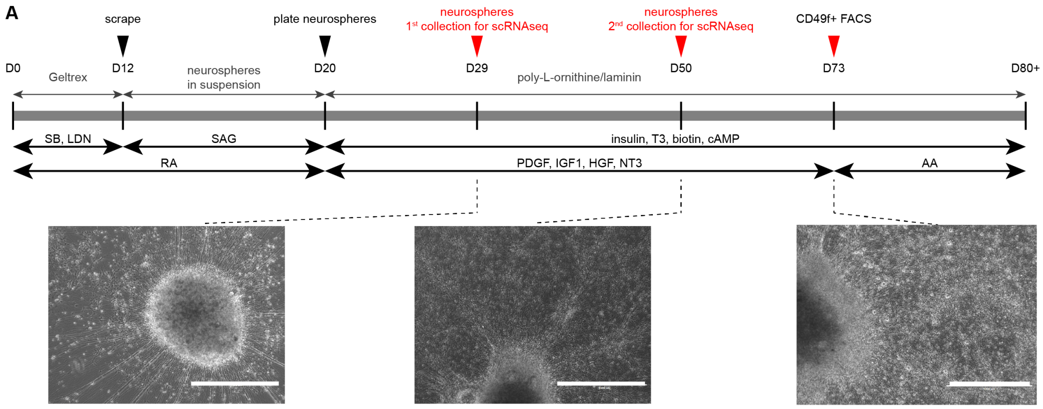


B

| | this study | | | | Barbar et al., 2020 | |
|---|---|---|---|---|---|---|
| | 30d #1 | 30d #2 | 50d #1 | 50d #2 | 73d (CD49f+) | 73d (unsorted) |
| preMUSCAT | 12,175 | 9,598 | 5,030 | 9,349 | 13,451 | 9,843 |
| postMUSCAT | 9,485 | 7,515 | 4,248 | 5,421 | 9,075 | 7,762 |
| % cells retained | 77.9% | 78.3% | 84.5% | 58.0% | 67.5% | 78.9% |

C D D

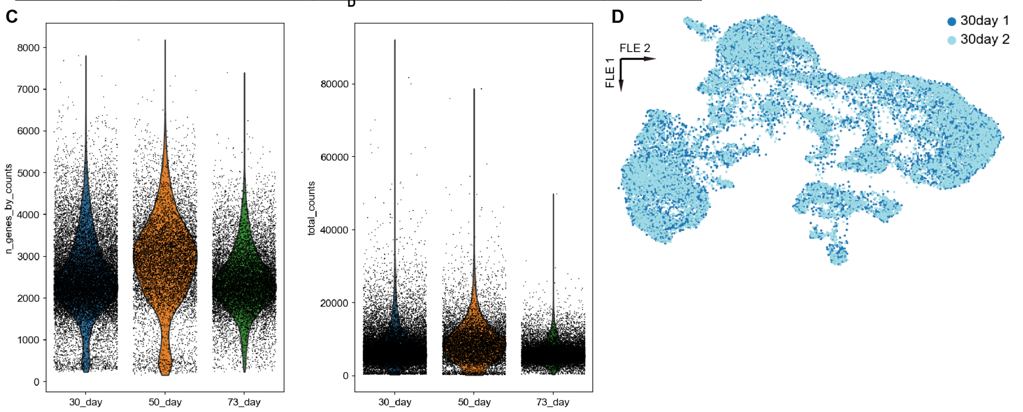


**Supplementary Figure 1 | Differentiation overview and scRNAseq analysis metrics. A)** Detailed human differentiation overview, with phase contrast micrographs of cells at the timepoints analyzed via scRNAseq. **B)** Table of quality control processing statistics for the newly produced data in this study and for the reanalyzed data from[36]. **C)** Number of genes identified for each cell and number of unique molecular identifier data for each cell. **D)** Harmony integration of two independently harvested and processed samples from the same human timepoint (30day) shows no evidence of batch effect. SB=SB431542, LDN=LDN193189; SAG= smoothened agonist; RA= retinoic acid; AA= ascorbic acid.

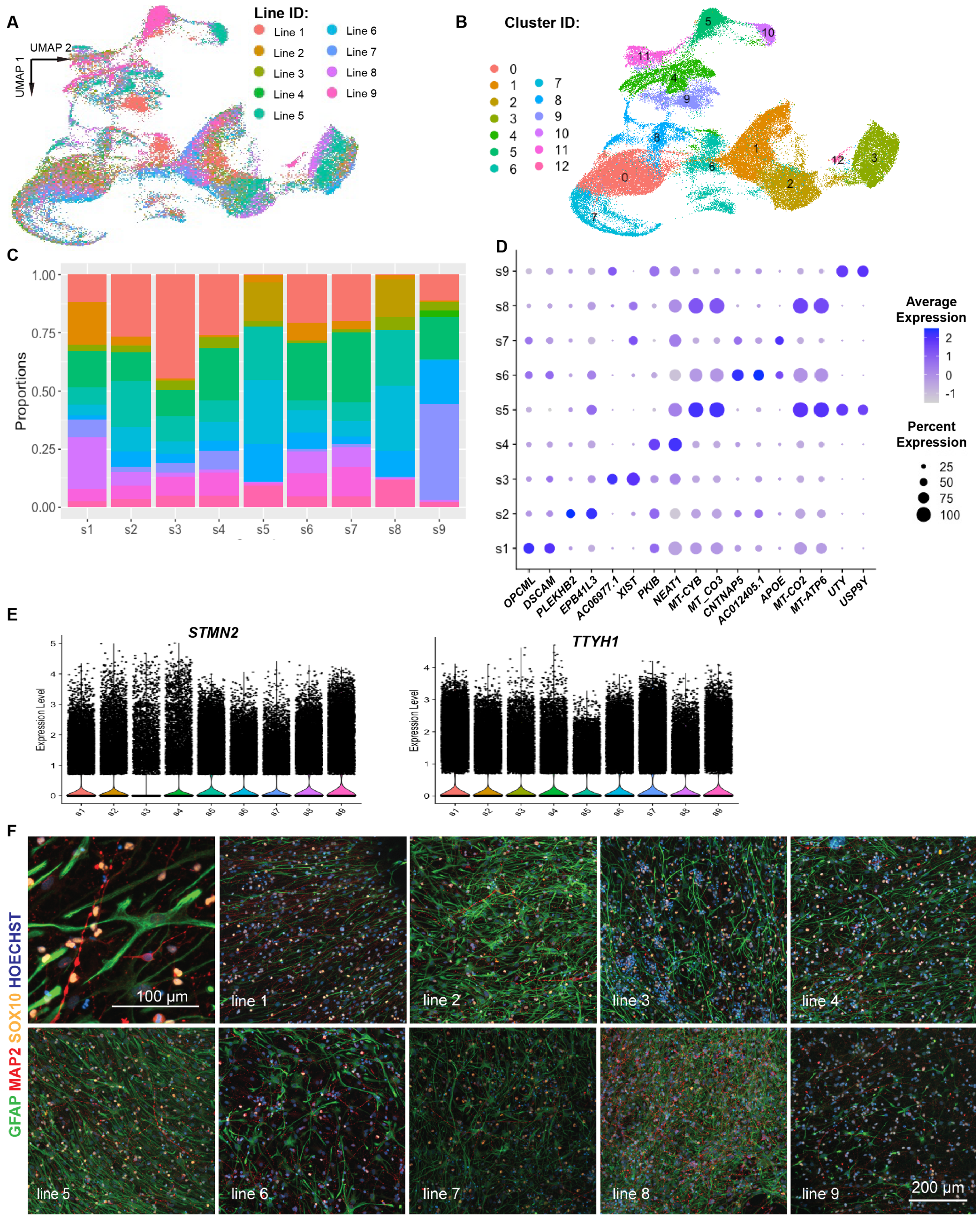


**Supplementary Figure 2 | Survey of differentiation cell type heterogeneity across 9 iPSC lines.** **A)** UMAP of 128,839 nuclei from 9 different iPSC lines (see Methods for line information). **B)** Clustering of nuclei from A) based on gene expression. **C)** Cell type proportion for each cluster identified in B) for each of the 9 iPSC lines tested suggest broad representation of each cell type in each iPSC line. **D)** Dot plot for the top 2 genes enriched in astrocytes from 9 different iPSC lines; differential expression analysis performed using the Wilcoxon ranked-sum test via Seurat. **E)** Normalized expression of *STMN2* and *TTYH1* shows similar expression of both genes across all 9 cell lines. **F)** One well of differentiated cells from each of the 9 lines were immunostained in parallel for cell type marker genes: GFAP (astrocytes; green), MAP2 (neurons; red), and SOX10 (oligodendrocytes; yellow) plus Hoescht 33342 nuclear stain (blue).

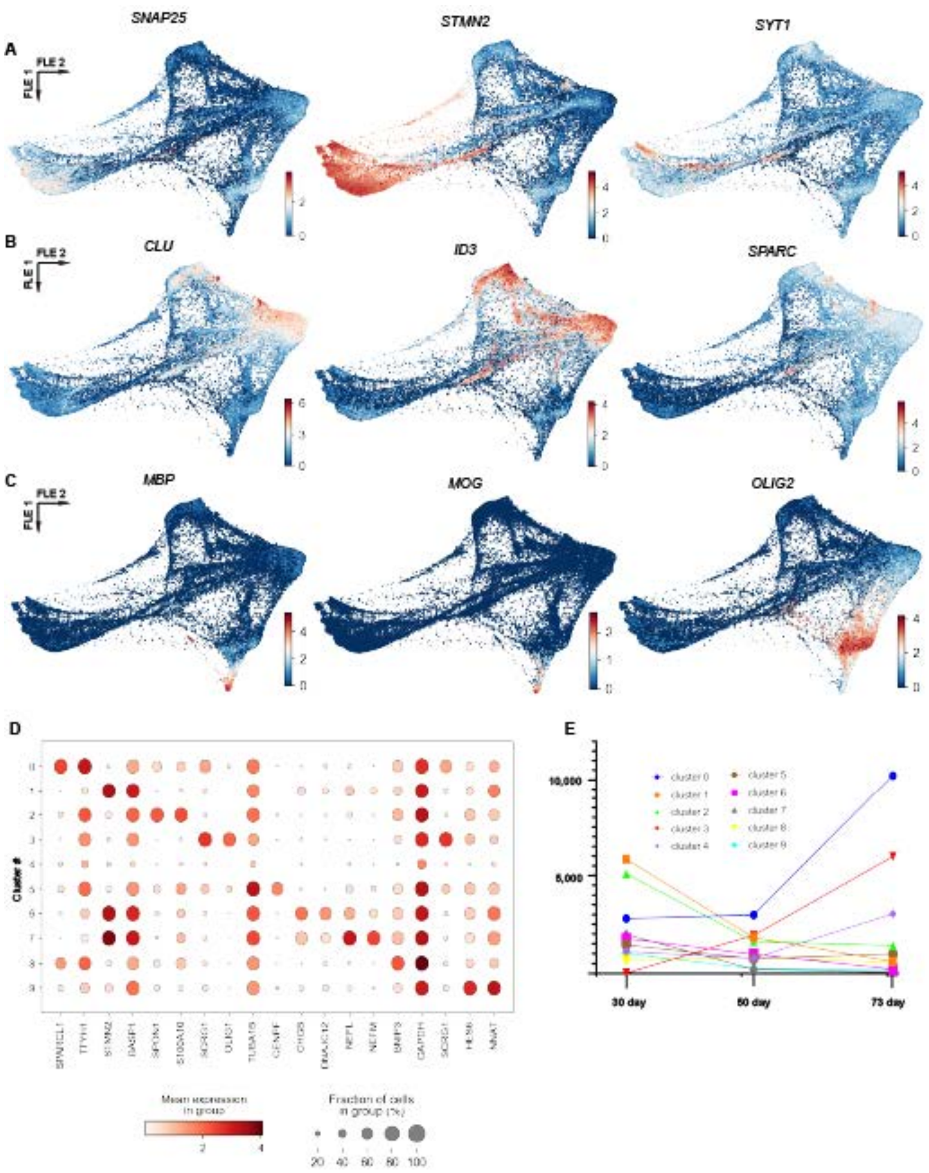


**Supplementary Figure 3 | Cell type feature plots. A)** Feature maps for neuron cell-type marker genes (*STMN2*, *SYT1*, *SNAP25*). **B)** Feature maps for astrocyte cell-type marker genes (*CLU*, *SPARC*, *ID3*). **C)** Feature maps for oligodendrocyte cell-type marker genes (*MOG*, *MBP*, *OLIG1*). **D)** Dot plot of top enriched genes for each cluster as determined by Scanpy (clusters correspond to those numbered in Fig 1D).

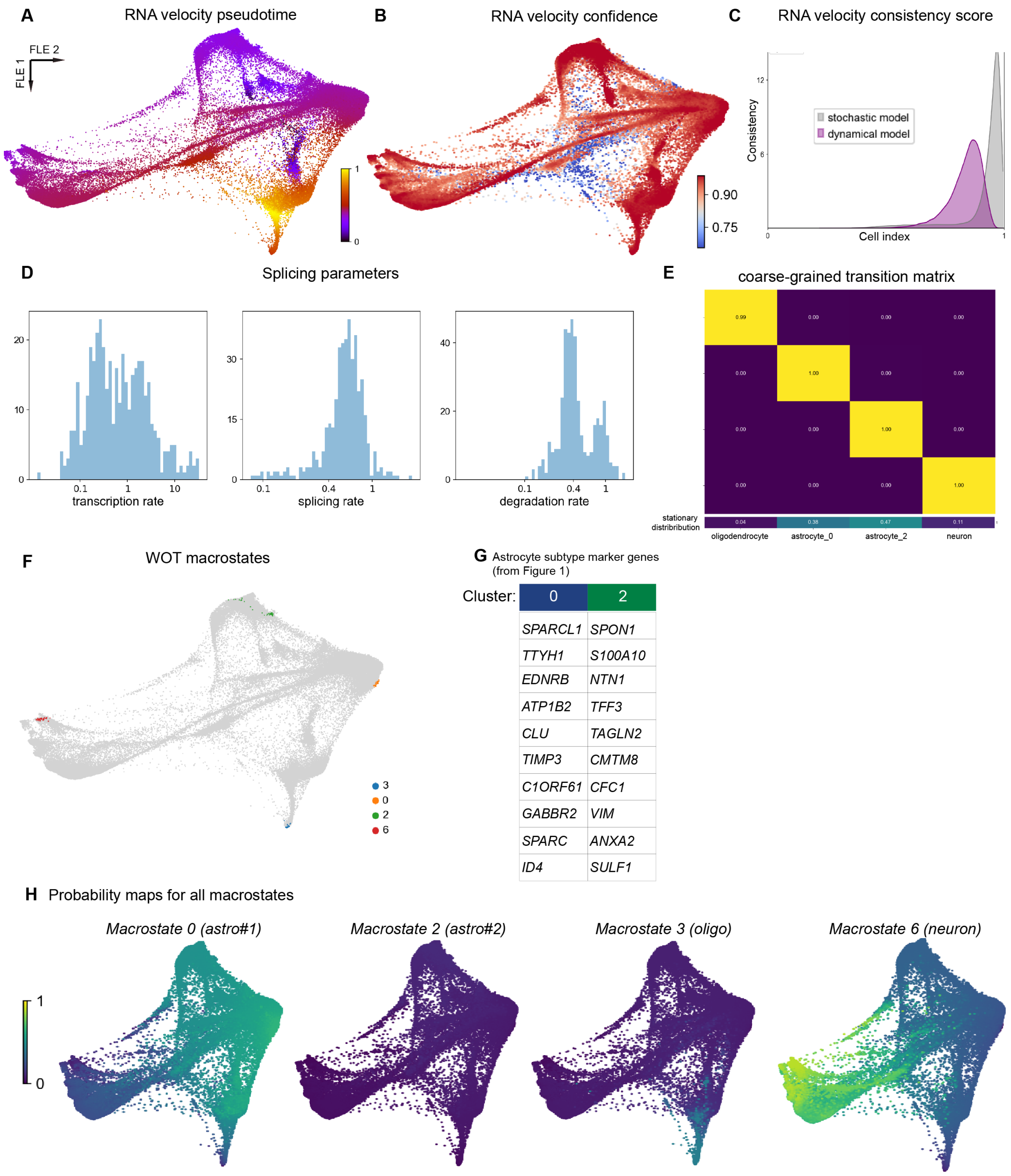


**Supplementary Figure 4 | RNA velocity and Waddington Optimal Transport (WOT) analysis. A)** Pseudotime calculated based on the stochastic model of RNA velocity. **B)** Confidence in RNA velocity calculated for each cell based on local coherence of velocity vectors[47]. **C)** Velocity consistency scores for each cell calculated for each of the two RNA velocity models employed above. **D)** Splicing parameters calculated using the dynamic model of RNA velocity. **E)** Plot of coarse-grained transition matrix depicts high probability to stay in one of the 4 macrostates upon entering the state (yellow boxes). **F)** Plot of four macrostates detected using the WOT algorithm. 0: astrocytes, 2: astrocytes, 3: oligodendrocytes, 6: neurons. **G)** List of marker genes for astrocyte cluster #2 from Fig 1D (cluster 2 in the original figure panel). **H)** Combined plot of probability maps for each macrostate graphed with same intensity scale. Replotting of data from Fig 1E-G and panel G from this figure.

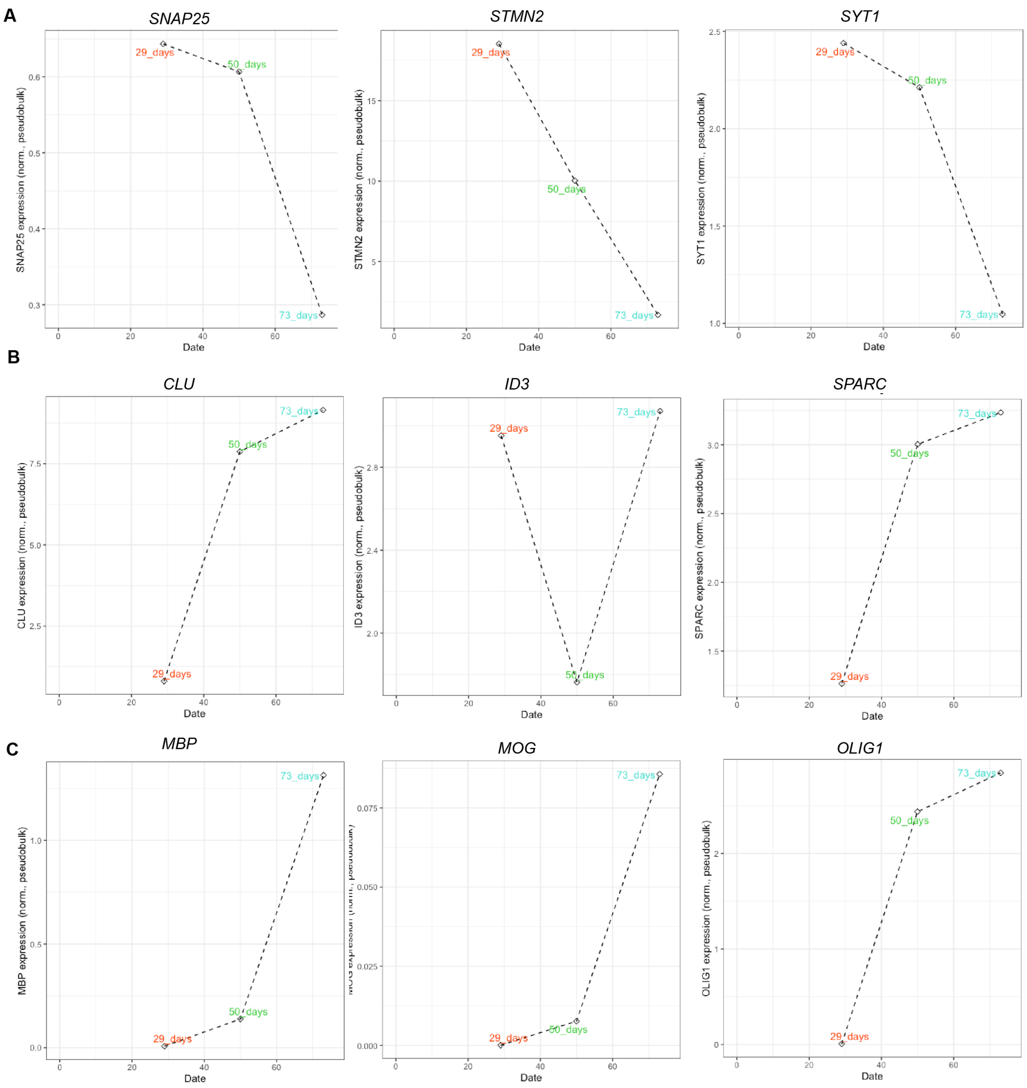


**Supplementary Figure 5 | Pseudobulk expression plots for human dataset cell-type marker genes. A)** Pseudobulk expression for neuron cell-type marker genes (*STMN2*, *SYT1*, *SNAP25*) at each timepoint shows peak expression during early timepoints, and decreasing expression over time. **B)** Feature maps for astrocyte cell-type marker genes (*CLU*, *SPARC*, *ID3*) show increasing expression over the course of the differentiation. **C)** Feature maps for oligodendrocyte cell-type marker genes (*MOG*, *MBP*, *OLIG1*) also show increasing expression over the differentiation, with particularly large jumps for *MOG* and *MBP* from day 50 to day 73.

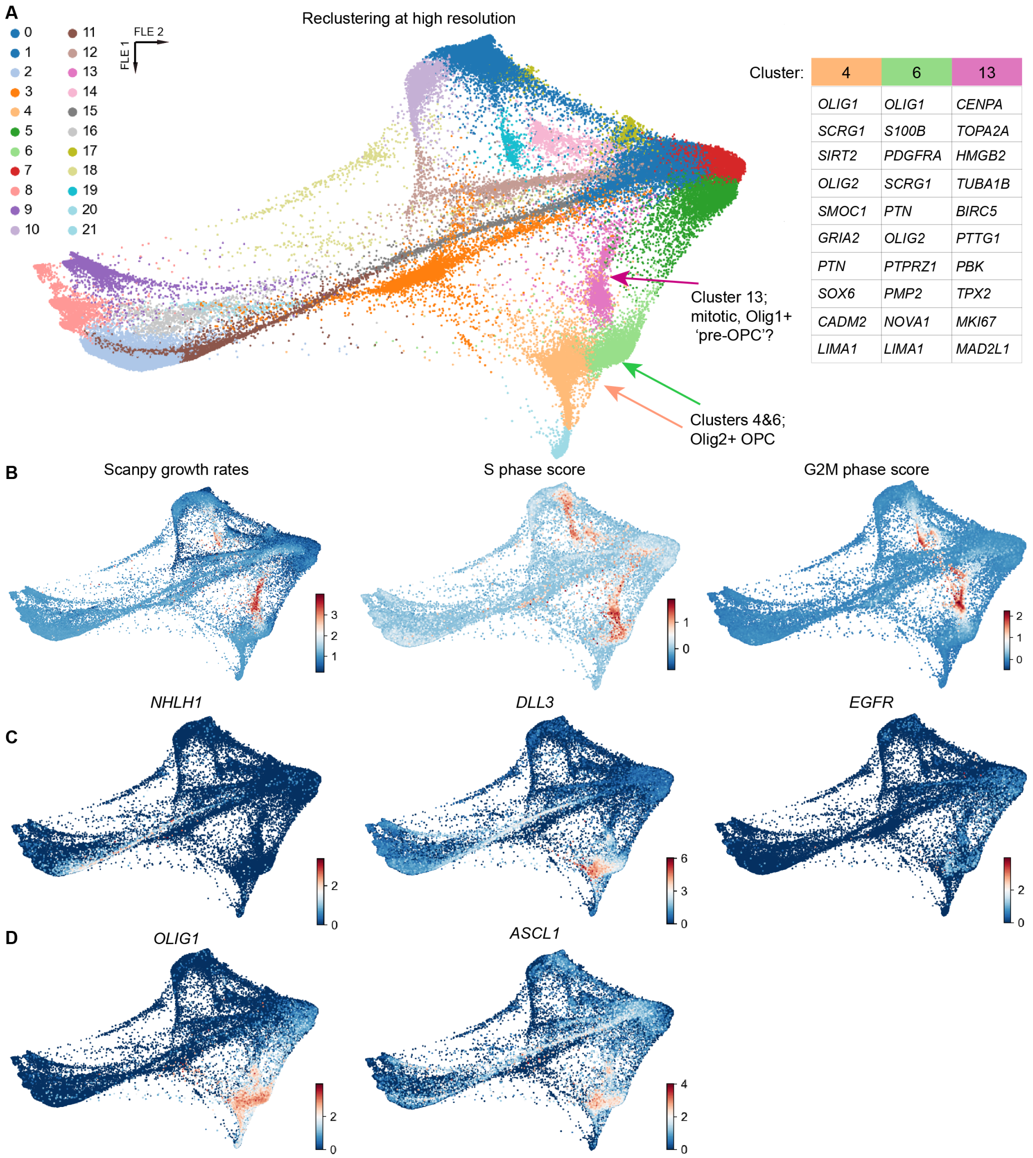


| 4 | 6 | 13 |
|---|---|---|
| *OLIG1* | *OLIG1* | *CENPA* |
| *SCRG1* | *S100B* | *TOPA2A* |
| *SIRT2* | *PDGFRA* | *HMGB2* |
| *OLIG2* | *SCRG1* | *TUBA1B* |
| *SMOC1* | *PTN* | *BIRC5* |
| *GRIA2* | *OLIG2* | *PTTG1* |
| *PTN* | *PTPRZ1* | *PBK* |
| *SOX6* | *PMP2* | *TPX2* |
| *CADM2* | *NOVA1* | *MKI67* |
| *LIMA1* | *LIMA1* | *MAD2L1* |

**Supplementary Figure 6 | Reclustering and analysis of transient states in human glial differentiation. A)** FLE reduction of the same data from Fig 1, reclustered at higher resolution (1.2). Marker genes for each of the clusters of interest are listed in the table to the right (see Results for further discussion), and the clusters are labeled with arrows on the FLE plot. **B)** Feature maps for mitotic scores from Scanpy87 ("Scanpy growth rates") and from scVelo[47] ("S phase score" and "G2M phase score"). **C)** Feature maps for genes enriched in shared neuron/astrocyte precursors (*NHLH1*, *DLL3*, *EGFR*). **D)** Feature maps for genes enriched in astrocyte/oligodendrocyte precursors (*OLIG2*, *ASCL1*).

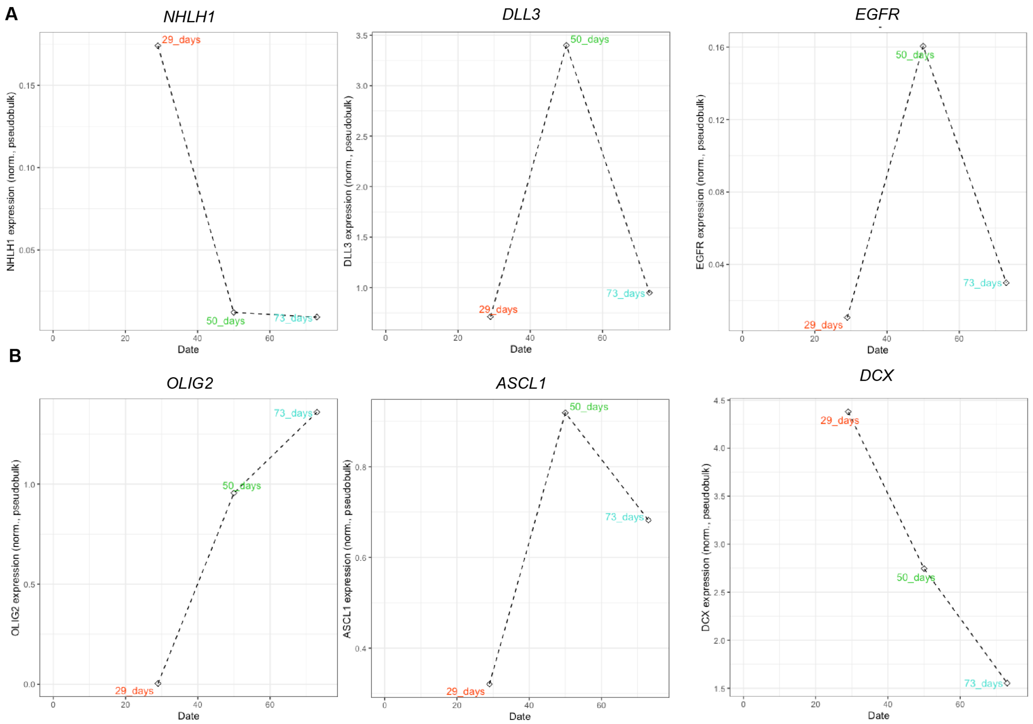


**Supplementary Figure 7 | Pseudobulk analysis of genes expression during transient states in human glial differentiation. A)** Pseudobulk expression for genes enriched in shared neuron/astrocyte precursors (*NHLH1*, *DLL3*, *EGFR*) shows an early peak of *NHLH1* expression, in contrast to *DLL3* and *EGFR* which peak during the intermediate 50 day timepoint. **B)** Pseudobulk expression for genes enriched in astrocyte/oligodendrocyte precursors (*OLIG2*, *ASCL1*) and in neural precursors (*DCX*).

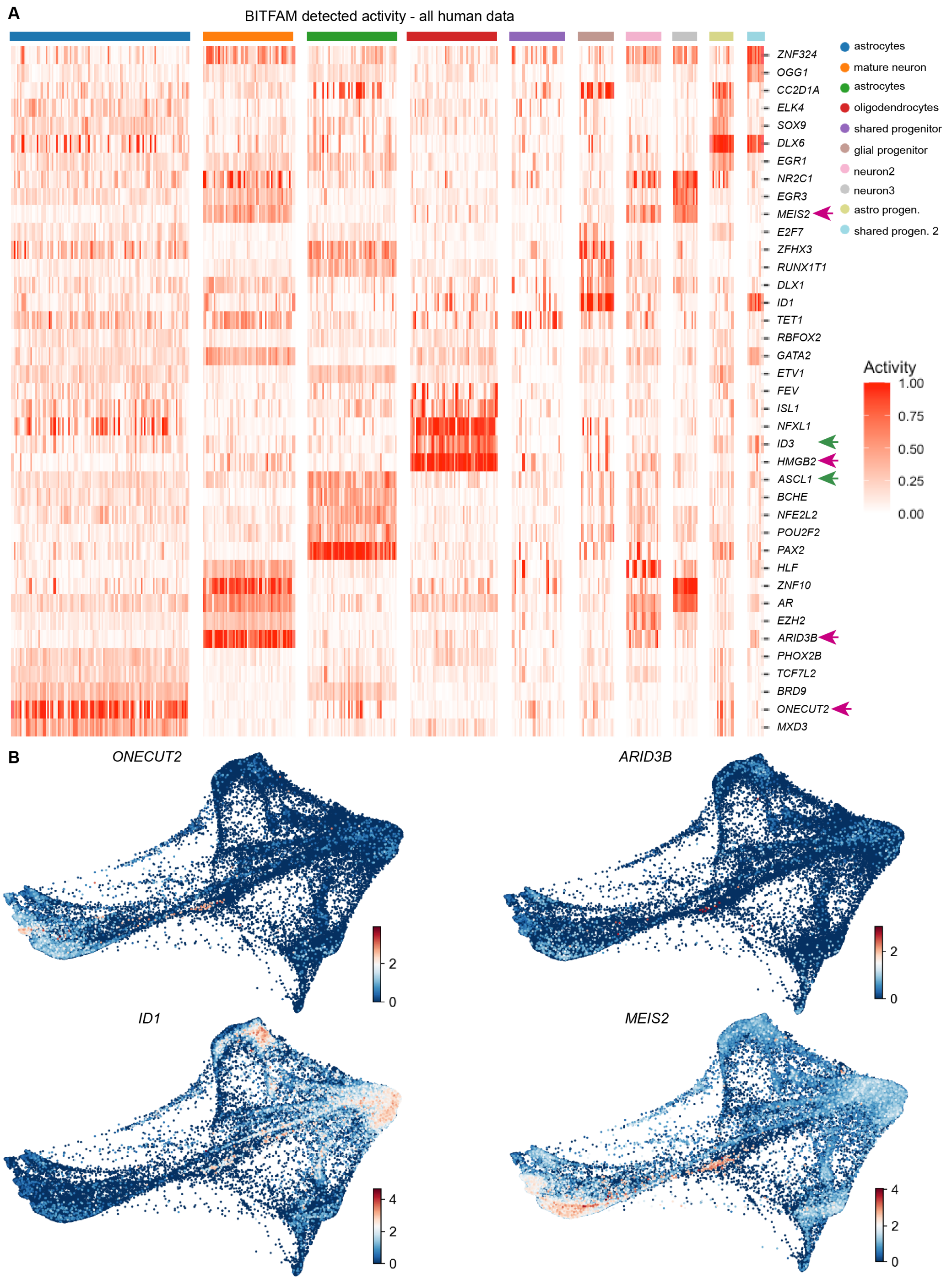


**Supplementary Figure 8 | Inferred transcription factor (TF) activity in differentiating human glia detected using BITFAM.** **A)** Heatmap of BITFAM detected activities for each cluster from Fig 1C (all human data timepoints). Clusters are numbered and labeled on the x-axis, and detected TFs are on the y-axis. Pink arrowheads: TFs with gene expression plotted below, green arrowheads: TFs with gene expression plotted in earlier supplemental figures. **B)** gene expression plots for genes highlighted with purple arrowheads in the BITFAM heatmap from A).

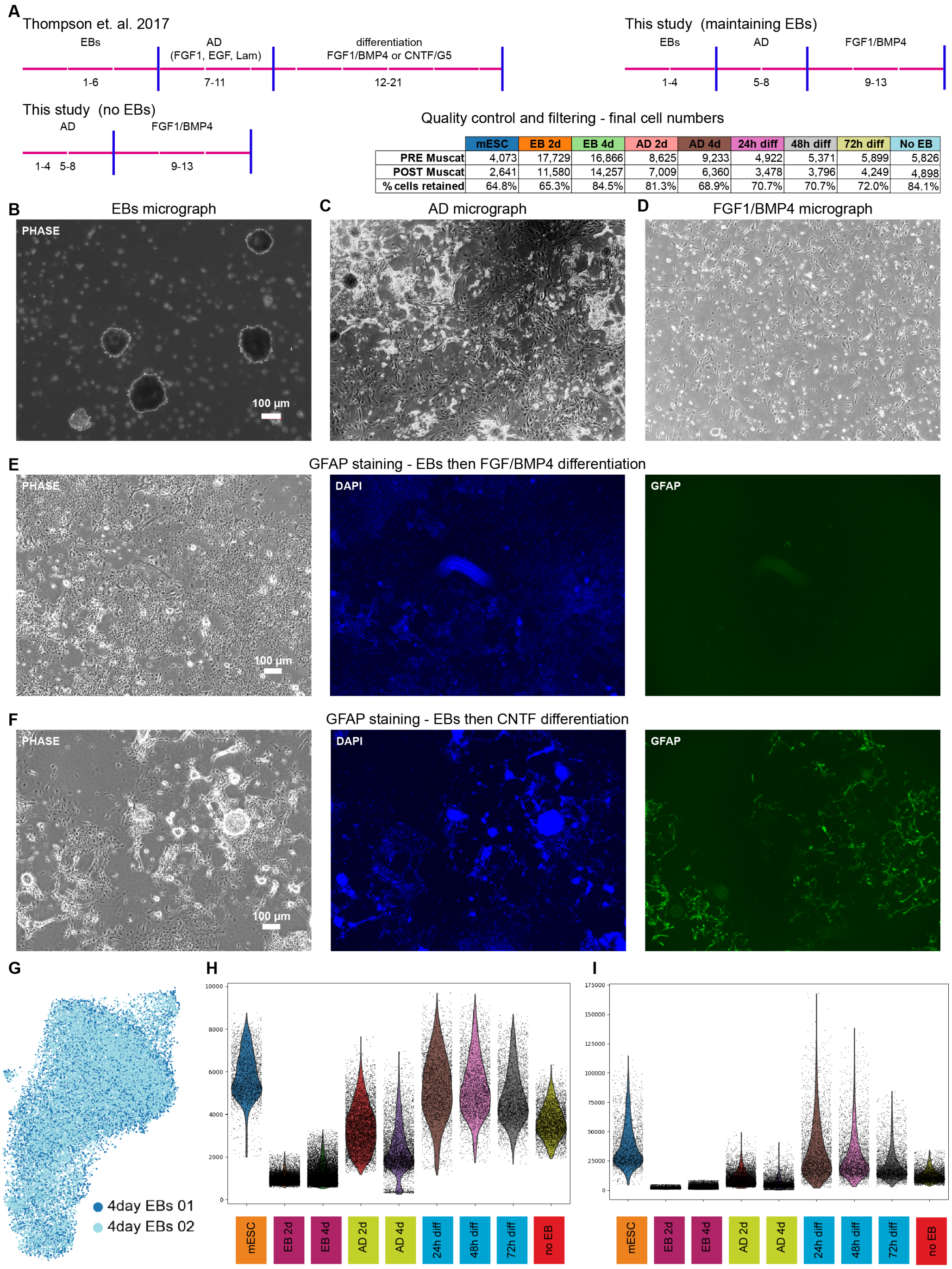


|  | mESC | EB 2d | EB 4d | AD 2d | AD 4d | 24h diff | 48h diff | 72h diff | No EB |
|---|---|---|---|---|---|---|---|---|---|
| PRE Muscat | 4,073 | 17,729 | 16,866 | 8,625 | 9,233 | 4,922 | 5,371 | 5,899 | 5,826 |
| POST Muscat | 2,641 | 11,580 | 14,257 | 7,009 | 6,360 | 3,478 | 3,796 | 4,249 | 4,898 |
| % cells retained | 64.8% | 65.3% | 84.5% | 81.3% | 68.9% | 70.7% | 70.7% | 72.0% | 84.1% |

**Supplementary Figure 9 | Differentiation overview and scRNAseq analysis metrics. A)** Mouse differentiation overview, with comparison across protocols. Quality control and pre/post filtering numbers for each mouse timepoint from the various protocols in table. **B-D)** Micrographs of cells during the embryoid body (EB), adherent differentiation (AD), and astrocyte growth factor stages. **E,F)** GFAP staining after protocol including EB stage. As expected based on published results[38], (E) minimal GFAP staining is visible following treatment with FGF and BMP4 but (F) robust staining is visible after CNTF treatment. **G)** Harmony integration of two independently harvested and processed samples from the same mouse timepoint (2 day EB) shows no evidence of batch effect. **H)** Number of genes identified for each cell and **I)** number of unique molecular identifier data for each cell from each timepoint.

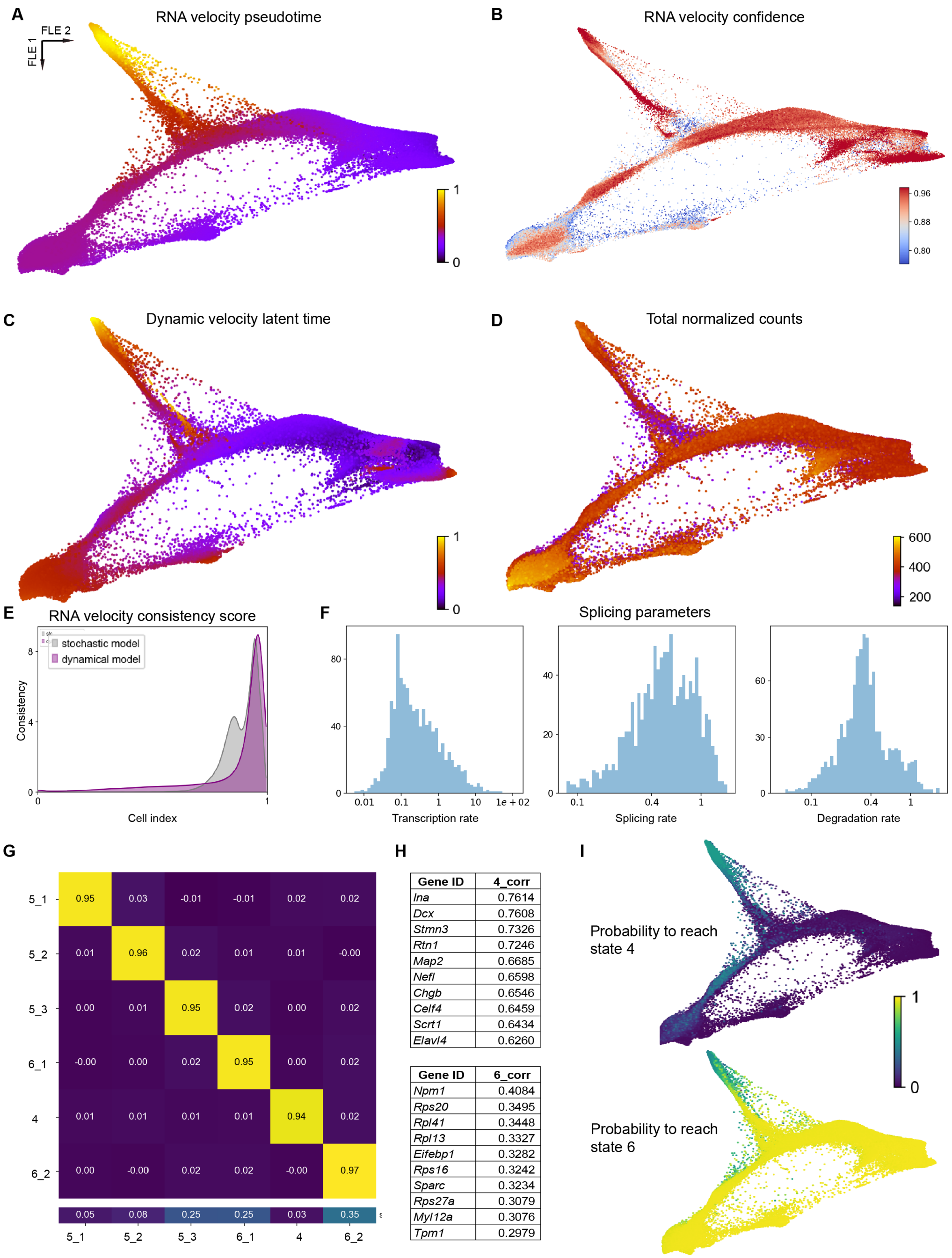


| Gene ID | 4_corr |
|---|---|
| *Ina* | 0.7614 |
| *Dcx* | 0.7608 |
| *Stmn3* | 0.7326 |
| *Rtn1* | 0.7246 |
| *Map2* | 0.6685 |
| *Nefl* | 0.6598 |
| *Chgb* | 0.6546 |
| *Celf4* | 0.6459 |
| *Scrt1* | 0.6434 |
| *Elavl4* | 0.6260 |

| Gene ID | 6_corr |
|---|---|
| *Npm1* | 0.4084 |
| *Rps20* | 0.3495 |
| *Rpl41* | 0.3448 |
| *Rpl13* | 0.3327 |
| *Eifebp1* | 0.3282 |
| *Rps16* | 0.3242 |
| *Sparc* | 0.3234 |
| *Rps27a* | 0.3079 |
| *Myl12a* | 0.3076 |
| *Tpm1* | 0.2979 |

**Supplementary Figure 10 | RNA velocity and Waddington Optimal Transport (WOT) analysis of mouse differentiation data. A)** Pseudotime calculated based on the stochastic model of RNA velocity. **B)** Confidence in RNA velocity calculated for each cell based on local coherence of velocity vectors. **C)** Latent time (similar to pseudotime) calculated based on the dynamic model of RNA velocity. **D)** Feature plot of normalized counts for each cell shows minimal differences across cells/timepoints. **E)** Consistency scores for each cell calculated for each of the two RNA velocity models employed above. **F)** Splicing parameters calculated using the dynamic model of RNA velocity. **G)** Plot of coarse-grained transition matrix depicts high probability to stay in one of the 2 macrostates upon entering the state (yellow boxes). **H)** Table of genes with highly correlated expression to enter one of the macrostates identified by WOT for each cluster. **I)** Combined plot of probability maps for each macrostate graphed with same intensity scale. Replotting of data from Fig 2F,G.

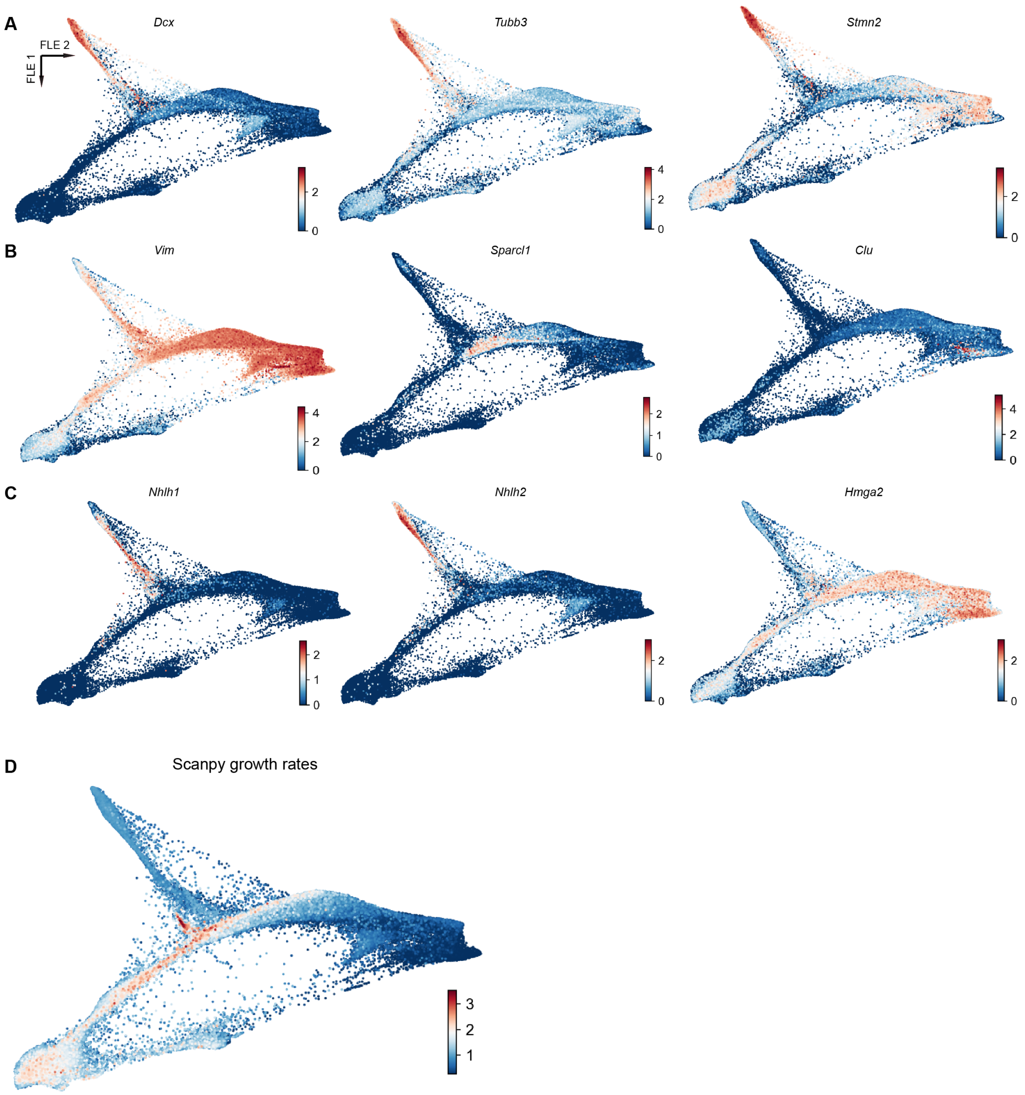


**Supplementary Figure 11 | Cell type feature plots. A)** Feature maps for neuron cell-type marker genes (*Dcx*, *Tubb3*, *Syt1*). **B)** Feature maps for astrocyte cell-type marker genes (*Vim*, *Sparcl1*, *Clu*). **C)** Feature maps for putative astrocyte precursor genes (*Nhlh1*, *Nhlh2*, *Hmga2*). **D)** Scanpy growth were calculated for each cell.

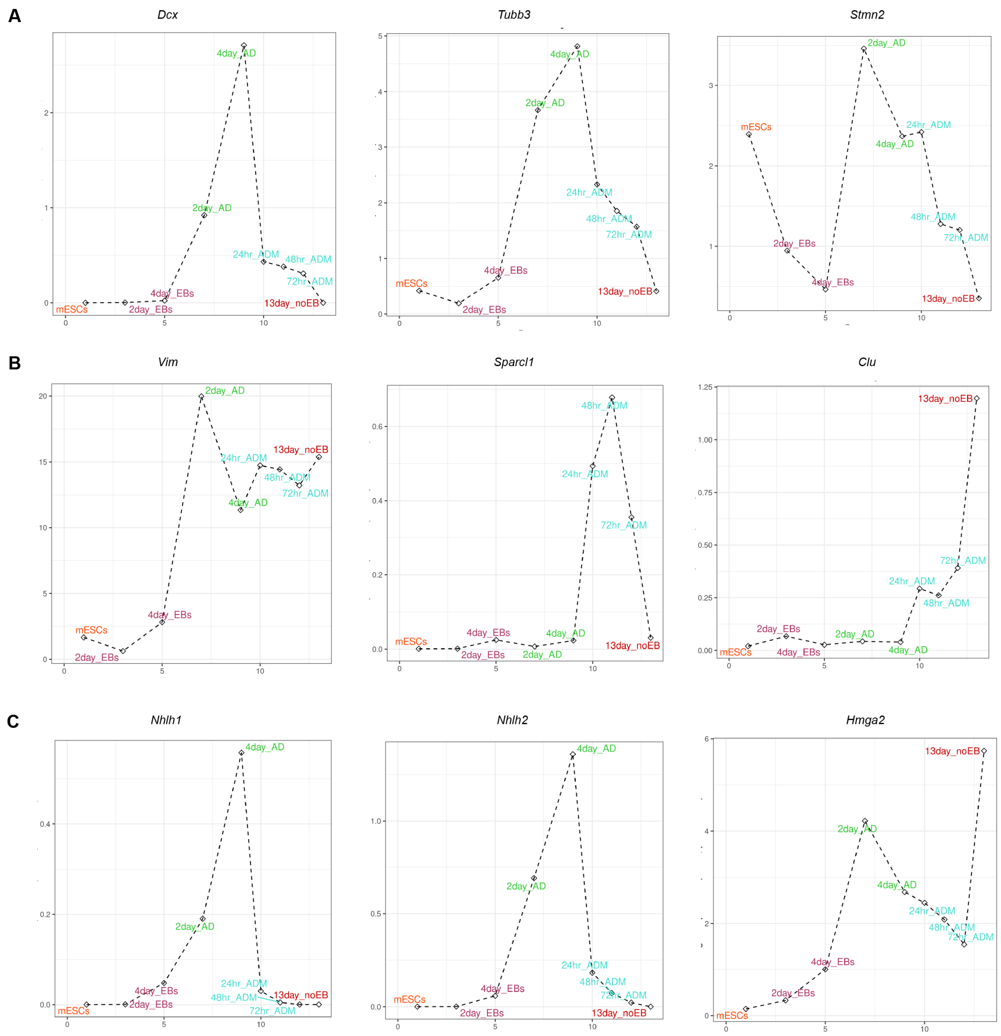


**Supplementary Figure 12 | Pseudobulk expression plots for mouse glial development.** **A)** Pseudobulk expression for neuron cell-type marker genes (*Dcx*, *Tubb3*, *Syt1*) at each timepoint shows a peak during intermediate stages of differentiation, then a decline during later stages. mESCs: mouse embryonic stem cells, EBs: embryoid bodies, AD: adherent differentiation, ADM: astrocyte differentiation media (FGF1+ BMP4). **B)** Pseudobulk expression for astrocyte cell-type marker genes (*Vim*, *Sparcl1*, *Clu*) show increased expression over time, with Vim increasing during the AD stage and the other markers increasing during the later ADM period. **C)** Pseudobulk expression for putative astrocyte precursor genes (*Nhlh1*, *Nhlh2*, *Hmga2*) show a similar pattern of increased expression during the AD stage, with a decrease during ADM.

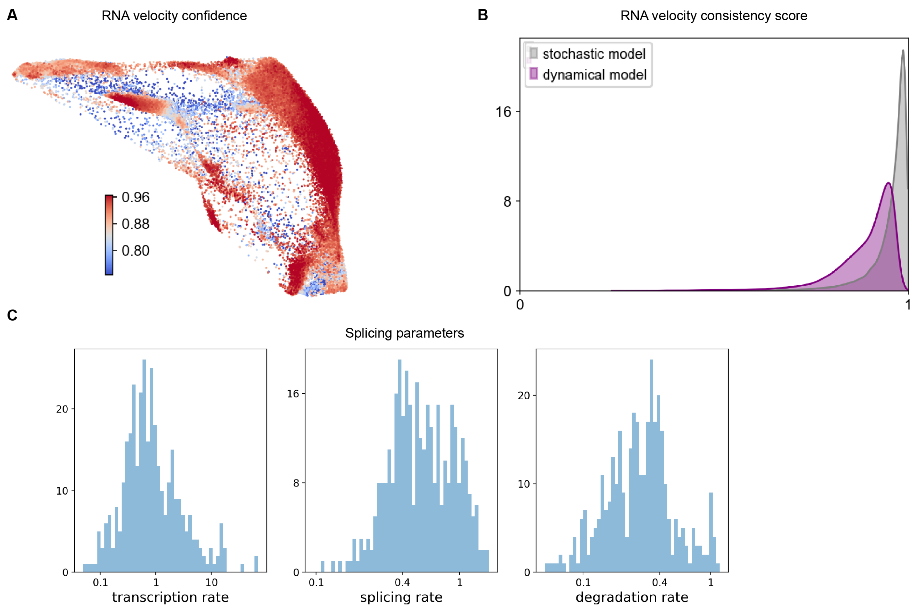


**Supplementary Figure 13 | RNA velocity analysis of subsets of mouse differentiation data.** **A)** RNA velocity confidence and **B)** RNA velocity consistency score for the subset of data shown in Fig 3 (adherent differentiation stages + growth factor stages). **C)** Splicing parameters calculated using the stochastic model of RNA velocity.

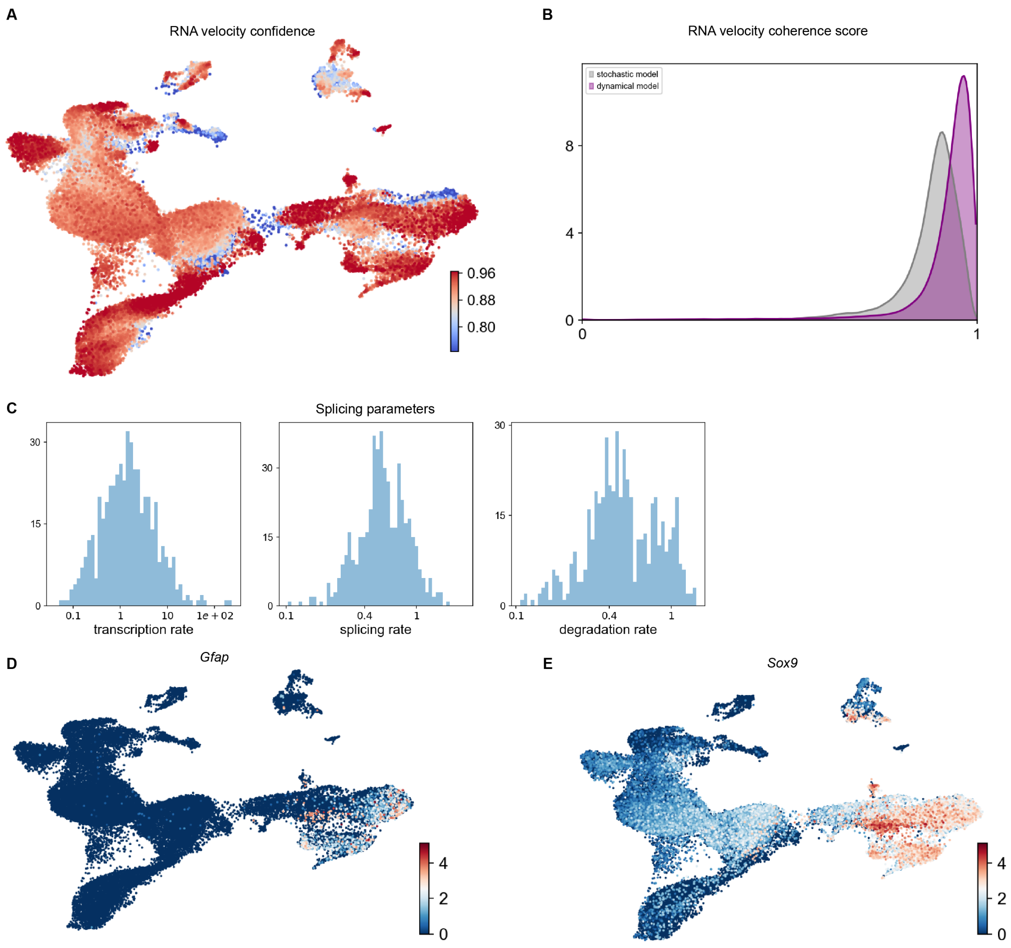


**Supplementary Figure 14 |** RNA velocity and Waddington Optimal Transport (WOT) analysis of subset mouse differentiation data integrated with previously published P3 primary mouse astrocyte data. **A)** RNA velocity confidence and **B)** RNA velocity consistency score for the subset of data shown in Fig 4 (adherent differentiation stages + FGF1/BMP4 stages, integrated with P3 primary astrocyte data). **C)** Splicing parameters calculated using the stochastic model of RNA velocity. **D)** Feature plot for Gfap shows lack of expression in differentiated cells, in addition to heterogeneous expression within the P3 data. **E)** Feature plot for Sox9 shows low expression in differentiated cells, but high albeit heterogenous expression in primary cells.

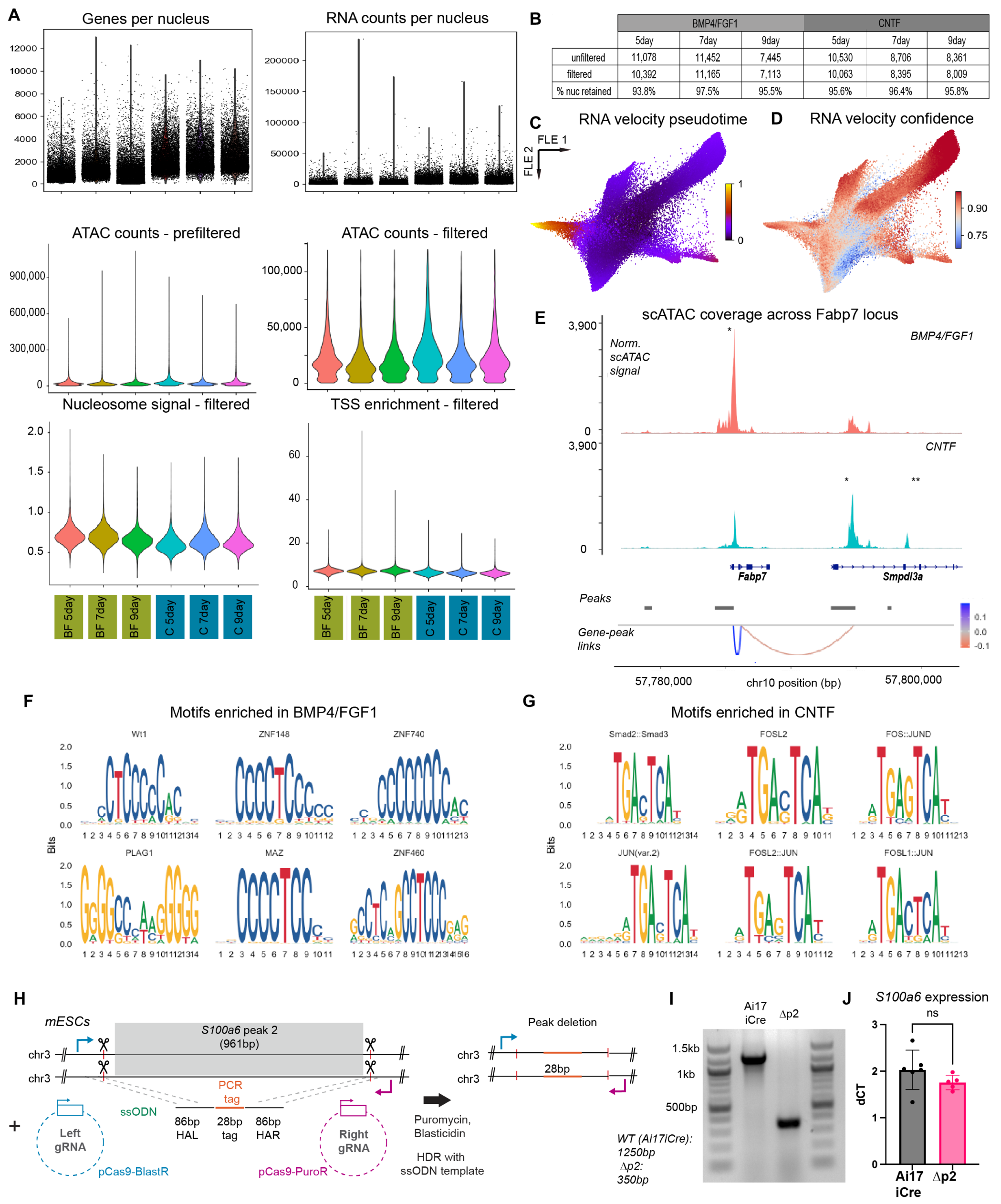


**Supplementary Figure 15 | Multiomics quality control and genomic accessibility analyses.** **A)** Violin plots of genes and counts per nucleus (top row), ATAC counts pre/post filter (middle row), and nucleosome signal and transcription start site enrichment scores (bottom row) for each timepoint. **B)** Table of each sample included in dataset analyzed for Fig 5. Nuclei were filtered based on the following cutoffs (see Methods): nCount_ATAC < 120000, nCount_ATAC > 500, nCount_RNA < 50000, nCount_RNA > 500, TSS.enrichment > 1. **C)** Pseudotime calculated based on the stochastic model of RNA velocity. **D)** Confidence in RNA velocity calculated for each cell based on local coherence of velocity vectors. **E)** Annotated coverage plot for *Fabp7* locus. See Fig 5F legend for detailed description of plot. **F,G)** Transcription factor motifs detected as enriched in peaks that were differentially accessible in BMP4/FGF1 (F) or CNTF (G) conditions. **H)** Approach for peak deletion for multiomic validation. **I)** Agarose gel demonstrating successful deletion of peak #2 and replacement with single-stranded oligo-donor nucleotide (ssODN) template. **J)** No differences in *S100a6* gene expression between the unedited (Ai17iCre) and the edited, peak #2 deleted (Δp2) mESC lines as measured by quantitative PCR.

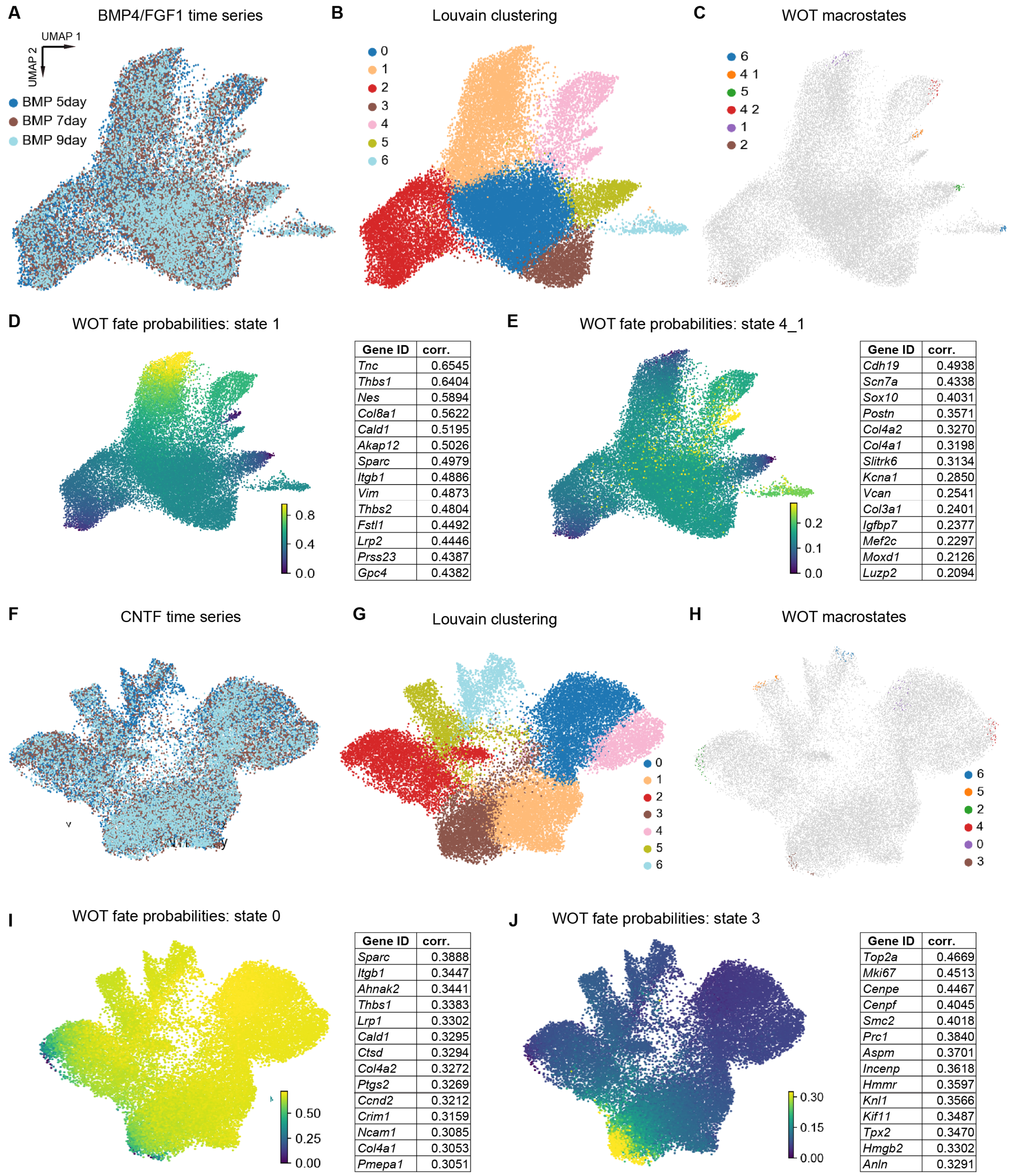


D)

| Gene ID | corr. |
|---|---|
| *Tnc* | 0.6545 |
| *Thbs1* | 0.6404 |
| *Nes* | 0.5894 |
| *Col8a1* | 0.5622 |
| *Cald1* | 0.5195 |
| *Akap12* | 0.5026 |
| *Sparc* | 0.4979 |
| *Itgb1* | 0.4886 |
| *Vim* | 0.4873 |
| *Thbs2* | 0.4804 |
| *Fstl1* | 0.4492 |
| *Lrp2* | 0.4446 |
| *Prss23* | 0.4387 |
| *Gpc4* | 0.4382 |

E)

| Gene ID | corr. |
|---|---|
| *Cdh19* | 0.4938 |
| *Scn7a* | 0.4338 |
| *Sox10* | 0.4031 |
| *Postn* | 0.3571 |
| *Col4a2* | 0.3270 |
| *Col4a1* | 0.3198 |
| *Slitrk6* | 0.3134 |
| *Kcna1* | 0.2850 |
| *Vcan* | 0.2541 |
| *Col3a1* | 0.2401 |
| *Igfbp7* | 0.2377 |
| *Mef2c* | 0.2297 |
| *Moxd1* | 0.2126 |
| *Luzp2* | 0.2094 |

I)

| Gene ID | corr. |
|---|---|
| *Sparc* | 0.3888 |
| *Itgb1* | 0.3447 |
| *Ahnak2* | 0.3441 |
| *Thbs1* | 0.3383 |
| *Lrp1* | 0.3302 |
| *Cald1* | 0.3295 |
| *Ctsd* | 0.3294 |
| *Col4a2* | 0.3272 |
| *Ptgs2* | 0.3269 |
| *Ccnd2* | 0.3212 |
| *Crim1* | 0.3159 |
| *Ncam1* | 0.3085 |
| *Col4a1* | 0.3053 |
| *Pmepa1* | 0.3051 |

J)

| Gene ID | corr. |
|---|---|
| *Top2a* | 0.4669 |
| *Mki67* | 0.4513 |
| *Cenpe* | 0.4467 |
| *Cenpf* | 0.4045 |
| *Smc2* | 0.4018 |
| *Prc1* | 0.3840 |
| *Aspm* | 0.3701 |
| *Incenp* | 0.3618 |
| *Hmmr* | 0.3597 |
| *Knl1* | 0.3566 |
| *Kif11* | 0.3487 |
| *Tpx2* | 0.3470 |
| *Hmgb2* | 0.3302 |
| *Anln* | 0.3291 |

**Supplementary Figure 16 |** Gene expression analysis of growth factor times series datasets in mouse ESC-differentiated astrocytes. **A-E)** Nuclei from the BMP4/FGF1 time series were plotted based on gene expression (A) and clustered via the Louvain algorithm at resolution 0.6 (B). The time series was analyzed via the Waddington Optimal Transport (WOT) algorithm (C), and fate probabilities were plotted for two WOT macrostates in (D) and (E). Genes correlated with WOT macrostate fate probability are plotted in tables next to their respective fate plots, see earlier figures and Methods for details. **F-J)** Nuclei from the CNTF time series were plotted based on gene expression (F) and clustered via the Louvain algorithm at resolution 0.6 (G). The time series was analyzed via the Waddington Optimal Transport (WOT) algorithm (H), and fate probabilities were plotted for two WOT macrostates in (I) and (J). Genes correlated with WOT macrostate fate probability are plotted in tables next to their respective fate plots.

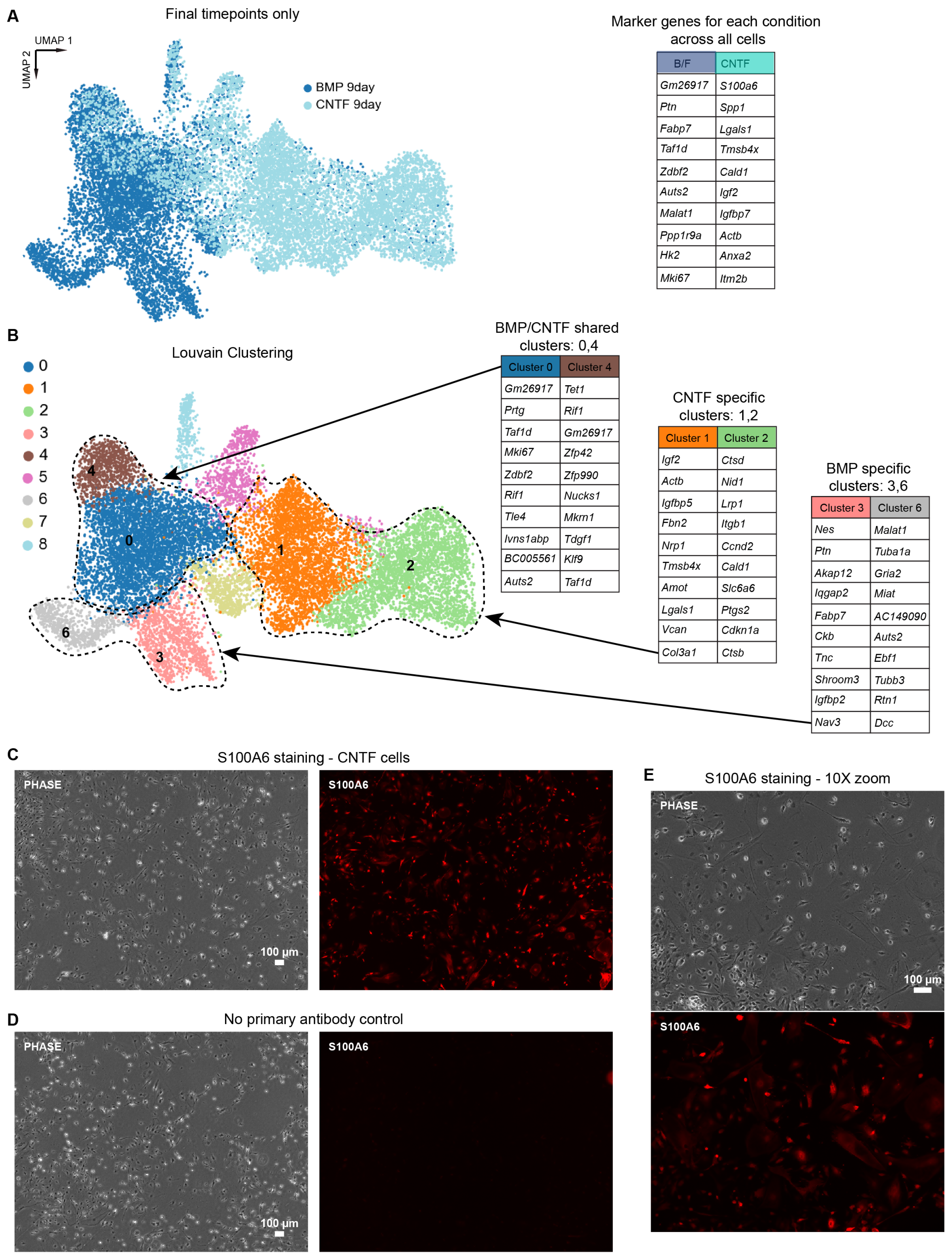


Marker genes for each condition across all cells

| B/F | CNTF |
|---|---|
| *Gm26917* | *S100a6* |
| *Ptn* | *Spp1* |
| *Fabp7* | *Lgals1* |
| *Taf1d* | *Tmsb4x* |
| *Zdbf2* | *Cald1* |
| *Auts2* | *Igf2* |
| *Malat1* | *Igfbp7* |
| *Ppp1r9a* | *Actb* |
| *Hk2* | *Anxa2* |
| *Mki67* | *Itm2b* |

BMP/CNTF shared clusters: 0,4

| Cluster 0 | Cluster 4 |
|---|---|
| *Gm26917* | *Tet1* |
| *Prtg* | *Rif1* |
| *Taf1d* | *Gm26917* |
| *Mki67* | *Zfp42* |
| *Zdbf2* | *Zfp990* |
| *Rif1* | *Nucks1* |
| *Tle4* | *Mkrn1* |
| *Ivns1abp* | *Tdgf1* |
| *BC005561* | *Klf9* |
| *Auts2* | *Taf1d* |

CNTF specific clusters: 1,2

| Cluster 1 | Cluster 2 |
|---|---|
| *Igf2* | *Ctsd* |
| *Actb* | *Nid1* |
| *Igfbp5* | *Lrp1* |
| *Fbn2* | *Itgb1* |
| *Nrp1* | *Ccnd2* |
| *Tmsb4x* | *Cald1* |
| *Amot* | *Slc6a6* |
| *Lgals1* | *Ptgs2* |
| *Vcan* | *Cdkn1a* |
| *Col3a1* | *Ctsb* |

BMP specific clusters: 3,6

| Cluster 3 | Cluster 6 |
|---|---|
| *Nes* | *Malat1* |
| *Ptn* | *Tuba1a* |
| *Akap12* | *Gria2* |
| *Iqgap2* | *Miat* |
| *Fabp7* | *AC149090* |
| *Ckb* | *Auts2* |
| *Tnc* | *Ebf1* |
| *Shroom3* | *Tubb3* |
| *Igfbp2* | *Rtn1* |
| *Nav3* | *Dcc* |

**Supplementary Figure 17 | Analysis of gene expression final timepoint from each growth factor time series. A)** (left) Dimensional reduction of gene expression data from final timepoints. (right) Top 10 genes positively enriched in cells belonging to one growth factor versus the other, as calculated in Scanpy. **B)** Louvain clustering of data from (A) identifies multiple clusters of cells based on gene expression that are either shared by both growth factor conditions (clusters 0,4), unique to BMP4/FGF1 condition (clusters 3,6), or unique to the CNTF condition (clusters 1,2). Top 10 marker genes for the above clusters are listed to the right. **C-E)** Validation of gene expression results with S100A6 immunostaining in cells from the final CNTF timepoint.

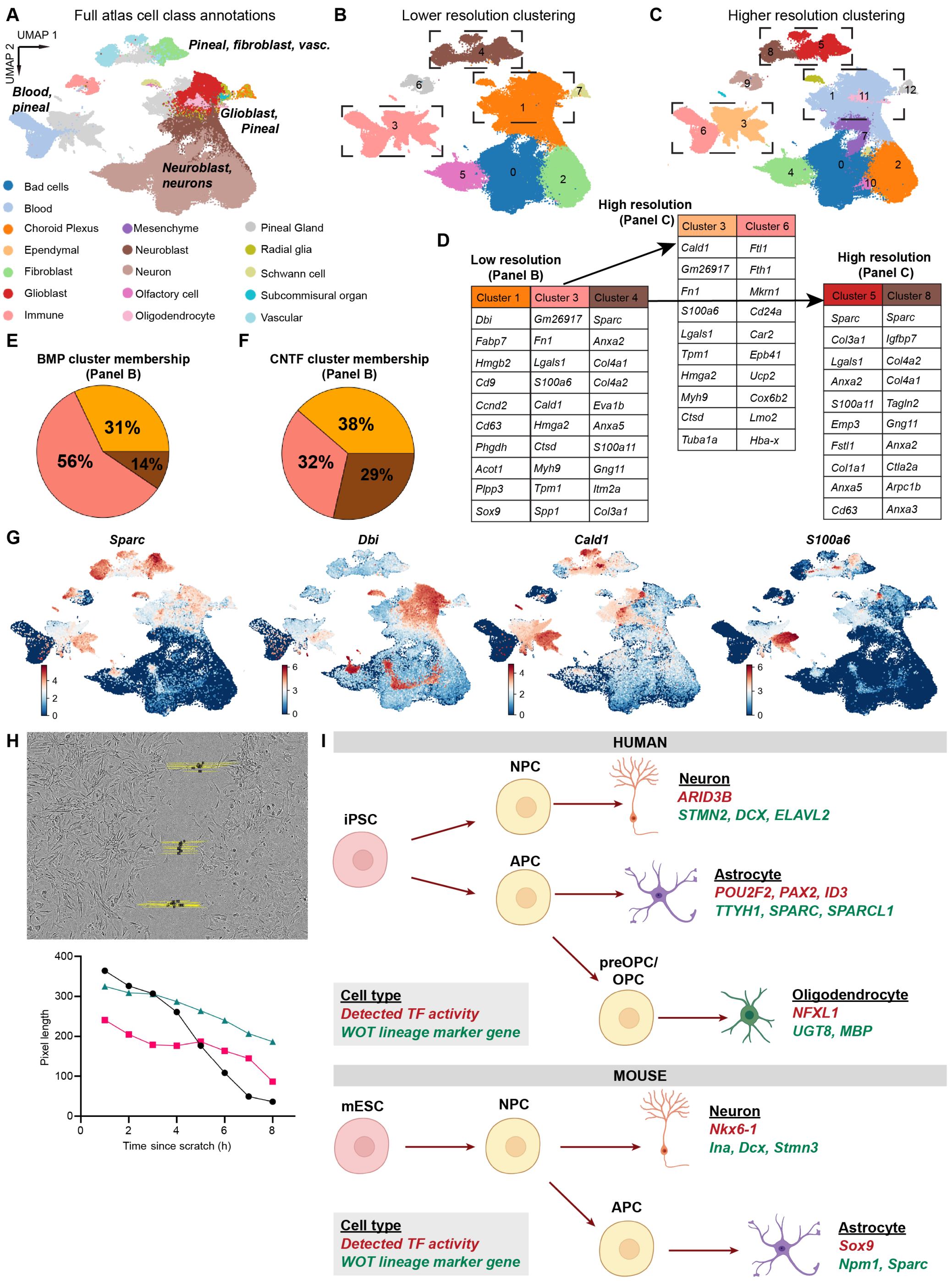


**Supplementary Figure 18 | Primary atlas integration, marker genes, and functional testing quality controls.** **A)** Cell class annotations were plotted from cell atlas metadata, and plotted for all 93,894 cells from the atlas. **B)** Cells were initially clustered at low resolution (0.1); reprint of Fig 6C for ease of comparison. **C)** Cells were clustered at higher resolution (0.3). **D)** Top marker genes for select clusters (dashed lines) from Fig S18B,C that contain both primary cells and differentiated cells. Marker genes for subclusters emerging from either Cluster 3 (left) or Cluster 4 (right) as part of re-clustering at higher resolution. **E,F)** Pie chart of percentage of cells from the BMP or CNTF 9 day timepoints (BMP: 7445 total, CNTF: 8361 total) that belong to each cluster from (B). **G)** Feature plots for major marker genes in panel (D). **H)** Still image from Movie S1 with example of scratch quantification (top). Graph of scratch closure over time from one well (bottom). **I)** Graphical summary of findings from the human datasets (top) and mouse datasets (bottom).
